# Molecular Mechanism of Nuclear Tau Accumulation and its Role in Protein Synthesis

**DOI:** 10.1101/2020.09.01.276782

**Authors:** Miguel Portillo, Ekaterina Eremenko, Shai Kaluski, Lior Onn, Daniel Stein, Zeev Slobodnik, Uwe Ueberham, Monica Einav, Martina K. Brückner, Thomas Arendt, Debra Toiber

## Abstract

Several neurodegenerative diseases present Tau accumulation as the main pathological marker. Tau post-translational modifications such as phosphorylation and acetylation are increased in neurodegenerative patients. Here, we show that Tau hyper-acetylation at residue 174 increases its own nuclear presence and is the result of DNA damage signaling or the lack of SIRT6, both causative of neurodegeneration. Tau-K174ac is deacetylated in the nucleus by SIRT6. However, lack of SIRT6 or chronic DNA damage result in nuclear Tau-K174ac accumulation. Once there, it induces global changes in gene expression affecting protein translation, synthesis and energy production. Tau-K174Q expressing cells showed changes in the nucleolus increasing their intensity and number, as well as in rRNA synthesis leading to an increase in protein translation and ATP reduction. Concomitantly, AD patients showed increased Nucleolin and a decrease in SIRT6 levels. AD patients present increased levels of nuclear Tau, particularly Tau-K174ac. Our results suggest that increased Tau-K174ac in AD patients is the result of DNA damage signaling and SIRT6 depletion. We propose that Tau-K174ac toxicity is due to its increased stability, nuclear accumulation and nucleolar dysfunction.

## Introduction

Neurodegeneration is characterized by the loss of brain function, including memory and learning impairments. Most neurodegenerative diseases are age-related, in fact ageing is the main risk factor for its development. One of the driving forces of ageing is DNA damage which has a profound effect on the brain, from its development to neurodegenerative phenotypes (1). Recently, it has been shown that Sirtuin family members play important roles in preventing ageing and age-related diseases (2–4). Among them, SIRT6 was shown to have a protective effect against the development of neurodegenerative phenotypes (5,6). SIRT6 is severely reduced in AD patients (6) and the lack of SIRT6 in the mouse model for brain specific KO (brSIRT6KO) results in a neurodegenerative phenotype with increased DNA damage and the appearance of hyper-phosphorylated Tau. Interestingly, although each neurodegeneration has its own signature and symptoms, several of them have Tau as common denominator. The functions of Tau in the cells range from stabilizing neuronal microtubules allowing axonal outgrowth, cargo transport and cellular polarity (7), to nuclear functions such as preventing genomic instability, rRNA transcription and chromatin relaxation (8–10). The toxic effects of Tau increase in a sporadic manner with age, together with the appearance of post-translational modifications (PTMs), including hyper-phosphorylation and acetylation (7,11,12). Various residues can be acetylated in Tau. Some of them increase Tau stability and toxicity (13–20), while others reduce them (21). Tau acetylation at residue 174 was associated with Alzheimer’s disease, preceding the appearance of phosphorylated-Tau (p-Tau) (16,22) (13,14,16,20). Intriguingly, Tau acetylation at residue K174 resembles several of the brSIRT6KO phenotypes in mice and cells, such as increased Tau stability and toxicity, behavioral defects, learning impairments, and increased neuronal cell death which results in neurodegeneration (6,16). Interestingly, Tau-K174ac effects on neuronal pathology are not due to the formation of neuro fibrillary tangles as the case of other Tau PTM’s (16,23). We previously showed that there is an accumulation of Tau in SIRT6 deficient cells and animals, partially due to GSK3 phosphorylation at residue S199 (6). However, we hypothesize that SIRT6 may play another role in Tau stability and function. We believe that it may regulate Tau by directly deacetylating it in the nucleus.

In this work, we define the mechanism leading to Tau-K174ac toxicity through acetylation due to DNA damage signaling and its nuclear accumulation. Tau acetylation at residue K174 is induced upon DNA damage by increased levels of CBP (Tau-acetyltransferase (20)). This acetylation induces Tau increased stability and nuclear localization. In the nucleus, it is deacetylated by SIRT6, however, its constant presence results in changes in gene expression, nucleolar increased function, Nucleolin accumulation and changes in protein translation rates. It is important to notice that these results are parallel to those observed in AD patients with increased nucleolar proteins, reduced SIRT6 levels and increased nuclear Tau-K174ac accumulation.

## Materials and methods

### Cell cultures

All cells were cultured in DMEM, 4.5g/l glucose, supplemented with 10% fetal bovine serum, 1% Penicillin and Streptomycin cocktail, and 1% L-glutamine. Cells were cultured with 5% CO_2_ at 37°C.

### Plasmids

Flag sequence was introduced into the plasmid, mEm-MAPTau-N-10, which was a gift from Michael Davidson (Addgene # 54155) (Em-Tau); ApoAlert™. The quick site-directed mutagenesis was used to introduce the K174Q mutation into the Tau WT using the following primers:

Tau K174Q-Fw: CAC CAG GAT TCC AGC ACA AAC CCC GCC CGC TC
Tau K174Q-Rev: GAG CGG GCG GGG TTT GTG CTG GAA TCC TGG TG

Tau K174Q S199E was created, through Quick change site-directed mutagenesis on Tau S199E, using the previously described primers.

### NAD^+^ consumption assay

Purified SIRT6-Flag was incubated at 37°C for 3h with either Tau 174n or Tau 174 acetylated peptide (10μg each) or H3K56Ac (10μg) with 2.5mM NAD^+^ and HEPES buffer (50mM HEPES pH 7.5, 100mM KCl, 20mM MgCl_2_, 10% Glycerol). After incubation samples were supplemented with 1uM 1,3-Propanediol dehydrogenase (1,3-PD) and 170mM 1,3 Propanediol for an additional 3h incubation.

SIRT6 NAD^+^ consumption was assessed — by measuring its absorption at 340nm — through NADH levels produced by 1,3-PDase activity.

To monitor spontaneous NAD^+^ consumption in the presence of Tau 174n or Tau 174 acetylated peptide, the assay was conducted without SIRT6 and each treatment was normalized to its control.

### Human brain tissue

It was used brain tissue of 10 AD patients and 8 healthy controls without neurological or psychiatric illness (Human sample table). The AD diagnosis was performed according to the neuropathologic assessment of Alzheimer’s disease (24,25). Case recruitment, autopsy and data handling were performed in accordance with the convention of the Council of Europe on Human Rights and Biomedicine and had been approved by the responsible local Ethical Committee.

### Immunohistochemistry

Tissue blocks containing human temporal cortex were immersed in 4% paraformaldehyde in phosphate buffer (0.1 M; pH 7.4) at 4°C. Blocks were subsequently immersed in 15% sucrose in phosphate-buffered saline (PBS). Frozen coronal sections were cut (30μm). Free floating sections were incubated in Citrate buffer (pH6.0) at 95°C for 10 min, subsequently transferred to TBS (pH7.4) and incubated with 3% H2O2 in TBS for 30 min to quench endogenous peroxidase activity. After 45 min of blocking unspecific binding sites at RT with 2% BSA (Serva), 0.3% dry milk and 0.5% donkey serum (Dianova) in TBS pH 7.4, sections were incubated with rabbit anti-Tau (Acetyl Lys174) antibody overnight at room temperature. The rabbit antibody was detected with biotinylated donkey anti-rabbit IgG (Dianova, Hamburg, Germany; 1:3,000). Sections were further processed with Extravidin-peroxidase conjugate (Sigma, Taufkirchen, Germany; 1:2,000) and 0.04% 3,3’-diaminobenzidine (DAB; Sigma) /NiSO4 /0.015% H2O2 as chromogen. After mounting, sections were covered with Entellan (Merck, Darmstadt, Germany), examined and digitized using an Axiophot microscope (Zeiss, Oberkochen, Germany) equipped with an AxioCam HRC camera (Zeiss).

### Human Samples Table

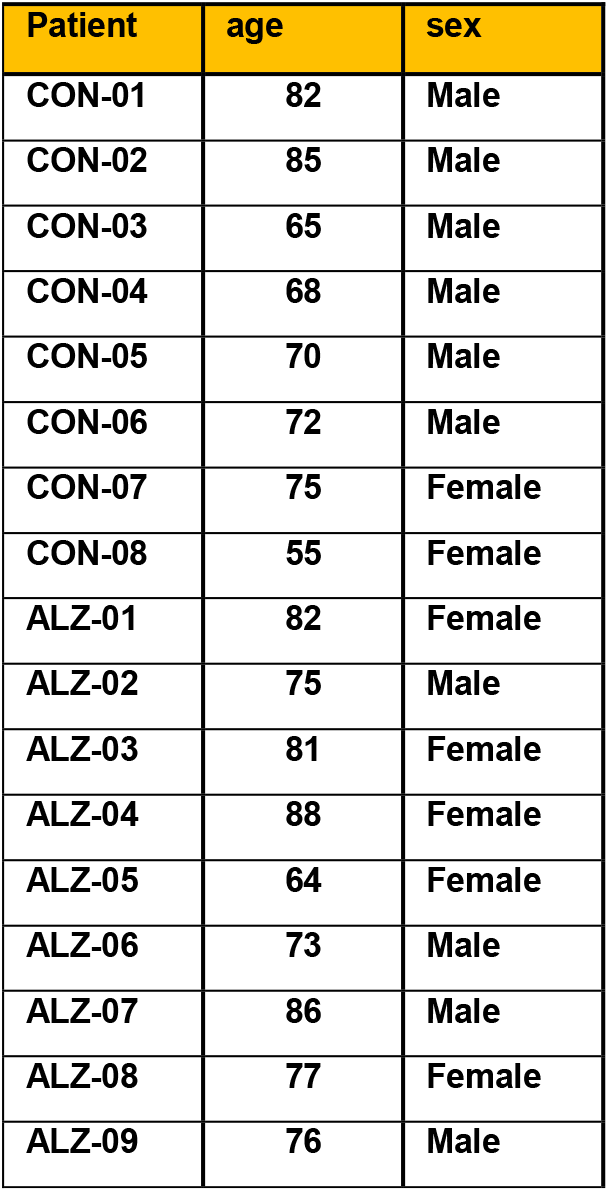

| Patient | age | sex |
| --- | --- | --- |
| CON-01 | 82 | Male |
| CON-02 | 85 | Male |
| CON-03 | 65 | Male |
| CON-04 | 68 | Male |
| CON-05 | 70 | Male |
| CON-06 | 72 | Male |
| CON-07 | 75 | Female |
| CON-08 | 55 | Female |
| ALZ-01 | 82 | Female |
| ALZ-02 | 75 | Male |
| ALZ-03 | 81 | Female |
| ALZ-04 | 88 | Female |
| ALZ-05 | 64 | Female |
| ALZ-06 | 73 | Male |
| ALZ-07 | 86 | Male |
| ALZ-08 | 77 | Female |
| ALZ-09 | 76 | Male |

### Immunofluorescence

SH-SY5Y cells were washed with PBS and fixed with 4% paraformaldehyde for 10 min at room temperature followed by two additional washes. Cells were permeabilized (0.1% NaCitrate, 0.1% Triton X-100, pH 6, in DDW) for 5 min and washed again. After 30 min blocking (0.5% BSA, 5% goat serum, 0.1% Tween-20 in PBS), cells were incubated with primary antibody diluted in blocking buffer over night at 4°C. The next day, cells were washed three times with wash buffer (0.25% BSA, 0.1% Tween-20 in PBS), incubated for 1h with secondary antibody (diluted in blocking buffer 1:200) at RT and washed three more times. Cells were then DAPI-stained for three minutes at RT and washed with PBS twice before imaging.

### RNA preparation

500,000 Cells were plated in 6-well plates for 4:30 hours, then transfected with 1ug of Tau mutant plasmids using PolyJet reagent (SL100688, SignaGen Laboratories, MD). Cells were collected 25 hours after transfection. RNA was then extracted using Nucleospin RNA Plus Kit (740984, MACHEREY-NAGEL GmbH & Co. KG, Duren, Germany).

### RNAseq

Libraries were prepared using QuantSeq 3’ mRNA-Seq Library Prep Kit FWD for Illumina (015.96, Lexogen GmbH, Vienna, Austria), and 400 ng of RNA were used per library. Samples were pooled (55 fmol cDNA per sample) and sequencing was performed using Nextseq5000 using 2 lanes. Sequenced used NextSeq 75 cycles (Illumina, Inc., CA).

12 samples were sequenced in the present study. These are composed of three biological replicates of SHSY5Y cells expressing Tau-WT and Tau-K174Q, Tau-S199E and Tau-S199E-K174Q. Sequencing resulted in an average of 8,239,157 reads per sample with an average sequence length of 86 bp. Raw reads were then trimmed for sequencing adaptors, poly-A, as well as poor-quality bases using Trim galore (v0.4.2; https://www.bioinformatics.babraham.ac.uk/projects/trim_galore/) and Cutadapt (v1.12.1; github.com/easybuilders/easybuildeasyconfigs/tree/master/easybuild/easyconfigs/c/cutadapt). After QC, an average of 8,197,527 reads per sample remained for downstream analysis with an average sequence length of 83 bp. Then, mapping and gene-level read count estimation were performed using STAR (v2.5.3a) and RSEM (v1.2.31), respectively, against the human reference genome (GRCh38). The DeSeq2 R package was used to normalize gene counts and perform differential gene expression, followed by clustering and visualizations using the DeSeq2 RLOG function (Variance Stabilizing Transformation). Clustering analysis was performed by employing a hierarchical clustering method (using Euclidean clustering metric and “ward.D2” agglomeration method), while the number of clusters were assessed using the “eclust” function from the “factoextra” R package. The “clusterprofiler” R package was used for Gene Ontology enrichment analysis, while gene annotation was retrieved from the ensemble biomart using the biomart R package. PCA was generated for the normalized count data using the plotPCA function from the DeSeq2 R package using the top 500 variable genes.

### Primary hippocampal culture

Primary hippocampal neurons were isolated from newborn C57BL/6 pups as previously described [40] with minor modifications. Briefly, hippocampi were collected into HBSS buffer, dissociated with papain (Sigma-Aldrich, P3125) and then seeded in chambers (μ-slide 4-well glass bottom, IBIDI GmbH Martinsried, Germany, 80427) coated with poly-L-lysine (Sigma-Aldrich, P-0899), using Neurobasal medium (GIBCO, 21103–049), supplemented with B-27 (GIBCO, 17504044), GlutaMAX (GIBCO, 35050–061) and 2% FBS (HyClone). After 24 h, the medium was replaced with a serum-free Neurobasal medium. After 10 days cells were infected with adenoviruses.

### Cloning of Tau WT and Tau K174Q into the adenoviral vector and virus preparation

FLAG-mEm-WT Tau and FLAG-mEm-Tau K174Q sequences were cloned into pAAV2 vector under human synapsin promoter (Addgene, #50465). Adenoviral particles were used to infect a primary hippocampal culture.

### SUnSET assay

SH-SY5Y neuroblastoma cells were plated and transfected the same day. Next day, prior to collection, cells were treated with 10ug/ml puromycin for 20 min. Protein extraction and immunoblot were performed using anti-puromycin specific antibody.

### 5-FU incorporation

Infected primary neurons expressing Tau WT and K174Q were treated with 2mM 5-Fluorouridine (5-FU) solved in DMSO for 15 minutes. Then fixation was performed with 2% paraformaldehyde during 10 min. Incorporation of 5-Fluorouridine was tracked using anti-BrdU (Sigma, B8434). Cellular segmentation and measurements were performed using Cell Profiler software, and the cells expressing Tau WT or K174Q were manually selected.

### CellTiter-Glo

SH-SY5Y cells were plated in a 96-well black coated with poly-L-ornithine. After plating, cells were transfected with 200 ng of Tau constructions. Next day, the plate was equilibrated at room temperature during half an hour and CellTiter-Glo^®^ Reagent (Promega, G7570) was added according to manufacturer instructions. The plate was mixed during 2 minutes on an orbital shaker and incubated at room temperature during 10 minutes prior to reading.

### Statistical analysis

Statistical analysis was done using GraphPad Prism 7. Analysis included either t-test or one-way ANOVA followed by a post-hoc Dunnet test or Tukey test, respectively. Significance was set at p<0.05.

Cellular fractionation and protein purification were performed as in Toiber et. al 2013

### Plasmids list

**Plasmids list:**
| Plasmid | Source | Cat. Number | PMID |
| --- | --- | --- | --- |
| CMV-Flag | Mostoslavsky Lab | - | 23911928 |
| mEmerald-MAPTau-N-10 Flag | Toiber Lab | - | - |
| mEmerald-MAPTau-N-10 FLAG K174Q | Toiber Lab | - | - |
| mEmerald-MAPTau-N-10 FLAG K174Q S199E | Toiber Lab | - | - |
| mEmerald-MAPTau-N-10 Flag S199E | Toiber Lab | - | - |
| pcDNA3β-Flag-CBP-HA | Addgene | 32908 | - |
| pQCXIP-GFP-LacR | Addgene | 59418 | 23929981 |

### Antibody list

**Antibody list:**
| Antibody | Company | Cat. Number | Method | Dilution |
| --- | --- | --- | --- | --- |
| Anti-Tau [E178] | Abcam | ab32057 | WB | 1:1000 |
| Tau 174Ac | Signalway Antibody | HW 181 | WB/IHC/IF | 1:1000/1:500 |
| βTubulin | Sigma-Aldrich | MAB1637 | WB | 1:1000 |
| HSC70 (B-6) | Santa Cruz | sc-7298 | WB | 1:1000 |
| Histone H3 | Abcam | ab1791 | WB | 1:3000 |
| Anti-Tau (phospho S199) | Abcam | ab4749 | WB | 1:1000 |
| SIRT6 | Abcam | ab88494 | WB | 1:1000 |
| H3K56ac [EPR996Y] | Abcam | ab76307 | WB | 1:1000 |
| CBP | Cell Signaling | D6C5 | WB/IF | 1:1000 |
| Anti-Actin Clone C4 | Merck Millipore | MAB 1501 | WB | 1:10000 |
| Phospho-Tau (Ser 404) | Cell Signaling | D2Z4G | WB | 1:1000 |
| Phospho-AMPKα (Thr172) (40H9) | Cell Signaling | 2535 | WB | 1:1000 |
| phospho-p70 S6 Kinase (Thr389) (1A5) | Cell Signaling | 9206 | WB | 1:1000 |
| Phospho-S6 Ribosomal Protein (Ser240/244) | Cell Signaling | 2215 | WB | 1:1000 |
| Nucleolin | Abcam | ab22758 | IF | 1:1000 |
| Fibrillarin [38F3] | Abcam | ab4566 | WB/IF | 1:2000/1:1000 |
| Anti-BrdU | Sigma-Aldrich | B8434 | IF | 1:1000 |
| Anti-Puromycin | DSHB | PMY-2A4 | WB | 1:1000 |
| Alexa Fluor 647 | Jackson ImmunoResearch |  | IF | 1:200 |
| Cross-Absorbed Secondary Antibody, Alexa Fluor 405 | ThermoFisher Scientific | A31556 | IF | 1:500 |

## Results

### SIRT6 regulates Tau stability through deacetylation at residue 174

We have previously showed that Tau levels are increased in SIRT6-KO (S6KO) cells and brSIRT6KO mice brains (Fig. S1A and B). Part of this increment is due to GSK3 activation and subsequent phosphorylation at Ser 199. However, S6KO cells expressing Tau-S199E point mutation (phospho-mimic Tau) still have higher levels of Tau (Fig. 1A), suggesting that its stability is likely to be affected by another SIRT6-dependent mechanism. Since SIRT6 is a lysine deacetylase, we hypothesized that SIRT6 could be regulating Tau deacetylation. To prove that, Tau K174-acetylated mimic (Tau-K174Q) levels were tested in S6KO cells. As previously described, Tau-K174Q showed increased stability in WT cells(16), but its effect was not considerably higher in S6KO cells suggesting that the increased stability is due to SIRT6 deficiency (Fig. 1B and C). Moreover, Tau-WT expression levels in S6KO were similar to those of acetyl mimic Tau, further supporting this hypothesis. T o detect Tau-174ac in our cell line we took advantage of the only available antibody aimed for this purpose whose specificity was tested first. Our results showed the suitability and specificity of this antibody for immunoblot and immunofluorescence (See Fig. S1C-E). Tau-K174ac levels are very low in the neuroblastoma cell line (see Fig. S1D), therefore we co-transfected cells with Tau-WT and CBP, a known Tau acetyltransferase (20,26). Tau hyper-acetylation at Lys 174 was increased by CBP expression and correlated with a slight increase in Total-Tau levels. Even without CBP, a significant increase was shown in S6KO cells, but this effect was accentuated by CBP overexpression (Fig. 1D-E, see 75 KDa band, monomeric form). These results suggest that Tau deacetylation at residue K174 required the presence of SI RT6. To test whether SI RT6 directly deacetylates Tau at residue K174, we purified Total Tau and performed an in vitro deacetylation assay on full length Tau (0N3R). To enrich Tau-K174ac we purified Total-Tau from HEK293T cells that were co-transfected with Tau-WT and CBP. Indeed, our results showed a decrease in Tau-K174ac signal and Tau fragmentation when incubated with SIRT6 in presence of NAD^+^ (Fig. 1F). Moreover, we used purified Tau-K174ac peptide and a non-acetylated peptide as control in a NAD^+^ consumption assay (NAD^+^ is required for SIRT6 activity, and it is hydrolyzed when SIRT6 is active, see Fig. S1F). As control, we measured NAD+ consumption when SIRT6 was incubated with a peptide sequence of its known target, H3K56ac (Fig. S1G). When SIRT6 was incubated with Tau-K174ac peptide, it consumed NAD+ resulting in its reduction. In contrast, this does not happen when incubated with the corresponding non-acetylated peptide (Tau 174n), thus confirming our results with full length Tau and supporting our claim that SIRT6 regulates Tau stability through deacetylation (Fig. 1G). We failed to observe the interaction of SIRT6 and Tau through regular IP, which could occur if that interaction was transient due its enzymatic nature, making it hard to detect. To circumvent this issue, we took advantage of Myc-SIRT6-BioID construct, which allows proximity-dependent biotinylation (see Fig. S1H). Tau levels are increased either by the absence of SIRT6 or after DNA damage, therefore, we hypothesized that DNA damage signaling could be a precondition for such interaction. Indeed, biotinylation of Tau was detected on irradiated cells but not in those which were not irradiated (Fig. 1H). These results suggest that SIRT6 and Tau interaction is DNA damage dependent, allowing the regulation of Tau by SIRT6. Overall, these results indicate that Tau is stabilized in SIRT6 deficient cells due to excess DNA damage and by increased acetylation levels at residue 174, and SIRT6 directly deacetylases Tau-K174ac (Fig. 1F).

**Figure 1.**
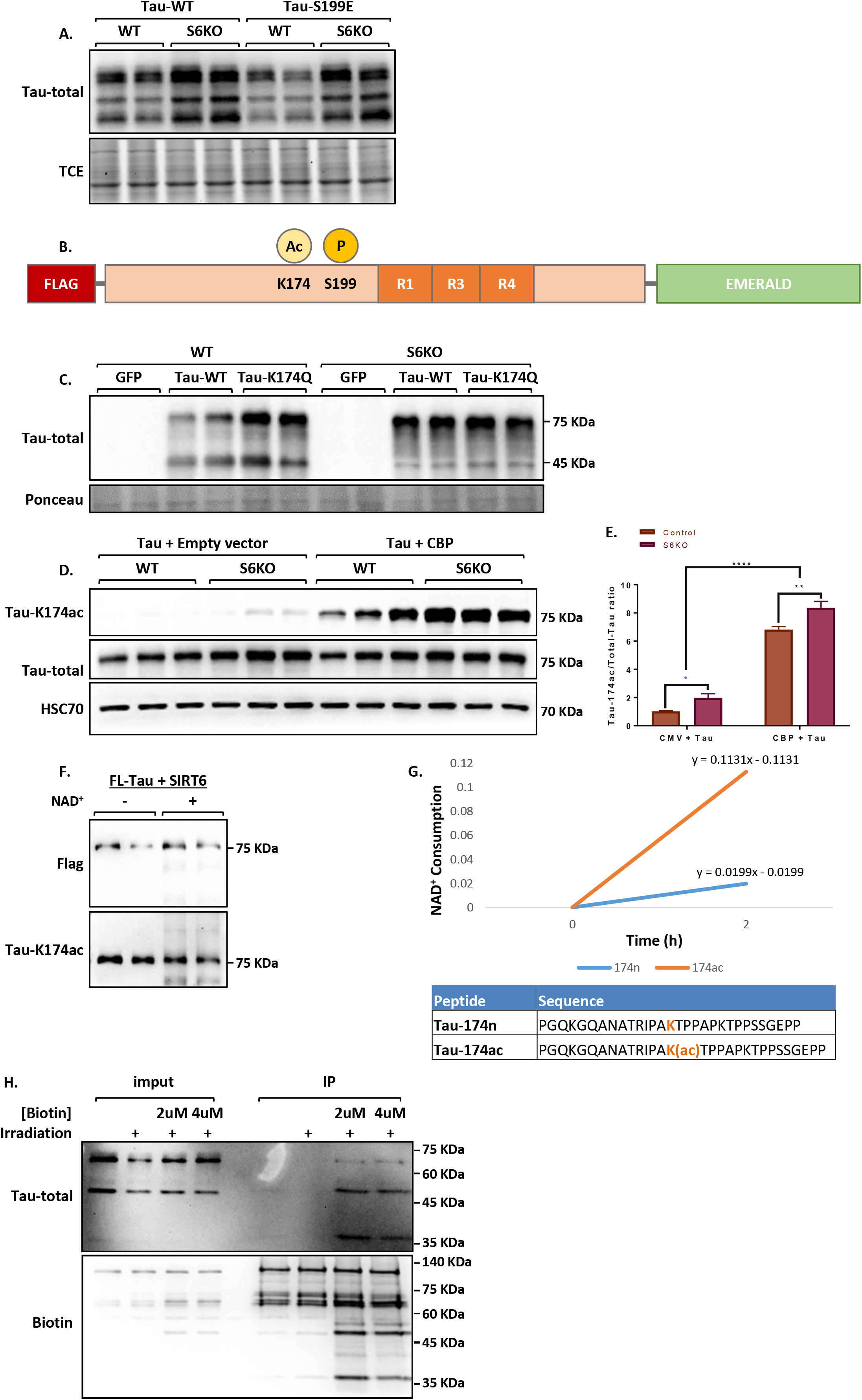
SIRT6 deacetylates Tau at residue 174. **(A)** Western blot of SHSY-5Y cells transfected with Tau-WT or Tau-199E. **(B)** Scheme of Tau 0N3R and its Acetylation at K-174, phosphorylation at S-199. **(C)** Western blot of WT and S6KO cells transfected with Tau-WT and Tau-K174Q (acetyl mimic) **(D)** Western blot of WT and S6KO cells co-transfected with Tau-WT with or without CBP. **(E)** Densitometry of Tau-174ac/Total-Tau from panel D. **(F)** *In vitro* deacetylation assay with SIRT6 and full-length Flag-Tau (Note that Tau was isolated from HEK293T cells that were co-transfected with CBP). **(G)** *In vitro* NAD+ consumption assay using acetylated (Tau-K174ac) and non-acetylated (Tau-174n) peptides, showing rates of conversion of remaining NAD+ to NADH in graph (1-(NADH levels)). **(H)** Proximity assay: Immunoprecipitation of biotinylated proteins performed on cells transfected with SIRT6-BioID irradiated with 4Gy. The quantification represents ANOVA analysis, and ttest * marked in blue showing +/- (SEM).

### Tau-K174ac is translocated to the nucleus

Since the subcellular location of SIRT6 is restricted to the nuclear compartment, the deacetylation of Tau by SIRT6 must occur there. First, we tested whether the increase of nuclear Tau was occurring in SIRT6 deficient brains (brS6KO), as well as in old mice and human AD patients. Indeed, in all of these samples Tau was significantly increased at chromatin fraction, confirming the critical role of SIRT6 in preventing its accumulation there (Fig. 2A-C). Moreover, chromatin fraction samples of AD patients showed an increase in nuclear Tau in general, as well as its acetylated and phosphorylated forms (Fig. 2C). Cytoplasmic fractions of AD brains also showed increased total Tau amounts, coupled with smearing (Fig. S2A). Next, we transfected Tau-WT or Tau-K174Q and measured Tau distribution in WT and S6KO cells by immunofluorescence. Mean intensity was measured at nuclear and cytoplasmic compartments, showing that Tau-WT is mainly cytoplasmic, however, in SIRT6 deficient cells the nuclear localization of Tau is increased. Interestingly, acetyl mimic Tau-K174Q levels were increased in the nucleus of both WT and KO cells (Fig. 2D-E), indicating that both the lack of SIRT6 or Tau acetyl-mimic result in an increase in nuclear Tau. We confirmed our results using protein fractionation to measure Tau levels in chromatin and cytoplasm, and the same phenomenon was observed (Fig. 2F). Lastly, we measured endogenous Tau-K174ac in the nuclear fraction of WT and S6KO cells, confirming the increase in Tau-K174ac in the absence of SIRT6, thus verifying SIRT6 regulatory function of this PTM (Fig. 2G). Remarkably, immunohistochemistry of control and AD patients’ cortex showed that in human brains, a specific antibody for Tau-K174ac detects its presence mainly in the nucleus. Moreover, in AD patients it was highly accumulated in the nuclear compartment (Fig. 2H and I, and S2B-E). Concomitantly, we also observed Tau-K174ac distribution in the nucleus in neuronal primary cultures infected with Tau-WT. Most of Tau-emerald localized to the cytoplasm, but Tau-K174ac was mainly nuclear, further supporting its preference for this cellular compartment (Fig. 2J). Overall, these results show that nuclear Tau is increased in aged mice brains or young SIRT6-deficient mice and in AD patients. Moreover, Tau acetylation at Lys 174 is an important residue for targeting Tau into the nuclear compartment.

**Figure 2.**
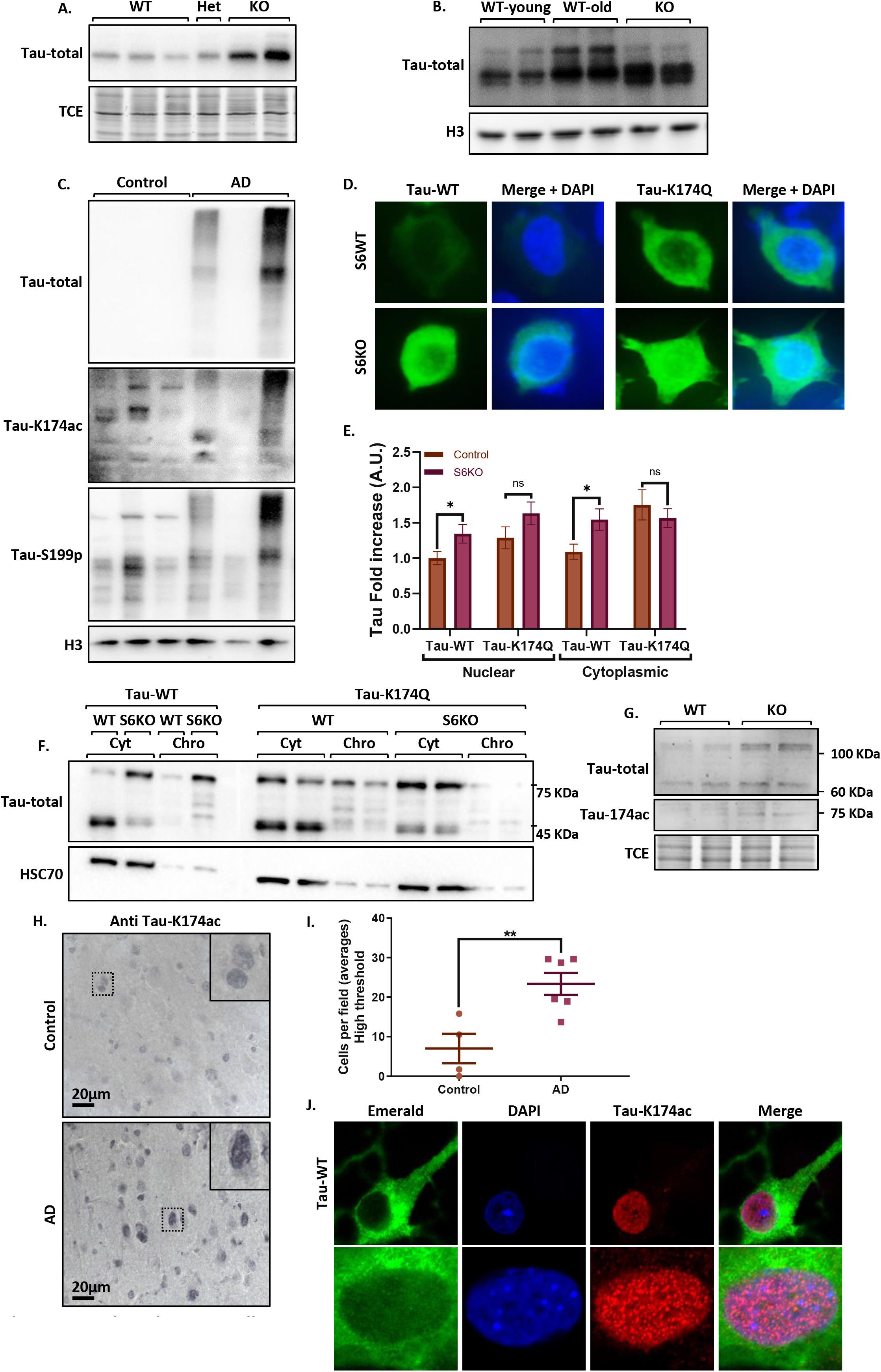
SIRT6 dependent Tau-Ac affect its location. **(A-C)** Western blots of chromatin fraction of proteins in: **(A)** mice brains from WT or S6KO mice, **(B)** young and aged mice (24m), **(C)** samples of brains from Alzheimer disease patients or nondemented controls **(D-E)** SHSY-5Y cells expressing mEmerald-Tau variants and the quantification of Tau at nuclear and cytoplasmic compartment+/- (SEM)--(n[WT+Tau-WT]=22, n[WT+Tau-K174Q]=25, n[S6KO+Tau-WT]=26, n[WT+TauK174Q]=21, p <0.05). **(F)** Western blot of chromatin and cytoplasmic fraction of transfected cells with Tau-WT and K174Q. **(G)** Western blot of nuclear fraction WT and S6KO cells transfected with Tau-WT. **(H)** Immunohistochemistry images of Control and AD brains with Tau174Ac **(I)** Number of Tau-174ac positive cells per field using high detection threshold. **(J)** Immunofluorescence of Tau-174ac neuronal primary culture infected with Tau-WT-mEmerald (upper panel with low exposure; lower panel with high exposure).

### Nuclear Tau increases upon DNA damage

DNA damage has a strong link to neurodegeneration. Thus, several genetic mutations in repair enzymes result in neurodegenerative symptoms. In addition, sporadic cases of AD, PD and ALS show an increase in DNA damage(1). SIRT6 deficient brains present an increase in genomic instability, therefore, we hypothesize that increased levels of nuclear Tau could be the result of DNA damage signaling. Tau was shown to be translocated into the nucleus under stress conditions such as heat shock and H2O2 treatments(27). Therefore, we hypothesized that nuclear Tau could also be induced by other stressful events, such as DNA damage. First, we tested whether DNA damage influences Tau nuclear translocation both in WT and S6KO cells. Indeed, Tau amounts at chromatin fraction increase upon DNA damage irradiation (I R) after 8hr from DNA damage stimuli (Fig. 3A). Strikingly, SIRT6 deficient cells, which cannot repair DNA properly and cannot deacetylate Tau-K174ac, presented higher levels of nuclear Tau at all times (Fig. 3A-B). To test whether Tau-174ac accumulates in the nucleus after DNA damage we irradiated primary neuronal cultures and neuron-like cells (SHSY 5Y), then we measured endogenous Tau-174ac mean intensity by immunofluorescence. As expected, Tau-174ac was located mainly at the nuclear compartment where its levels increased upon irradiation, further supporting our claim that either SIRT6 deficiency or DNA damage influence Tau permanence at the nucleus by means of acetylation at Lys 174 (Fig. 3-C-D and Sup. F3D-E). Interestingly, Tau 174ac accumulates at the nucleus forming foci (Tau foci), therefore we measured whether irradiation induces changes in their properties. Indeed, Tau foci number, their intensity and the area occupied in the nucleus increased 8 hours post-irradiation in both primary neuron culture and SHSY5Y cell line (Fig 3E and Sup. 3A-C and F-I), when SIRT6 levels are at their lowest (see Fig. 3A). This supports the role of SIRT6 in preventing Tau nuclear accumulation.

**Figure 3.**
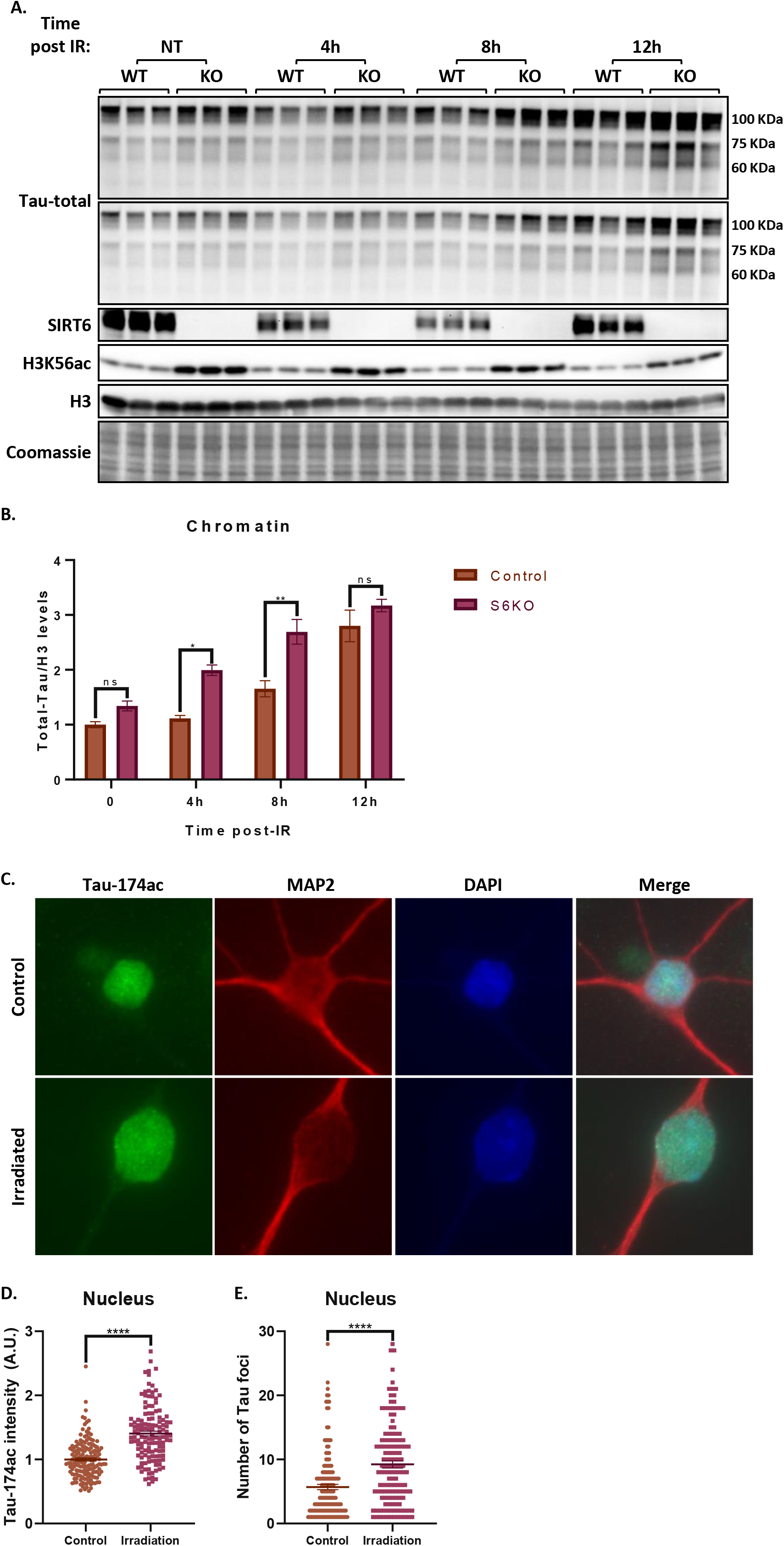
DNA damage induces Tau translocation to the nucleus. **(A)** Time lapse of nuclear Tau accumulation after irradiation of SHSY-5Y WT and S6 KO cells. **(B)** Densitometry of panel A **(C-E)** Immunofluorescence of Tau-174ac in primary neuronal culture, quantifications of three independent experiments. **(C)** Immunofluorescence of Tau-174ac performed in primary culture of brain -/+ irradiation. **(D)** Normalized Tau-174ac mean intensity at nucleus for -/+ Irradiation. **(E)** Number of Tau foci at nucleus -/+ Irradiation. Error bars represent (SEM) (n[-Irradiation] =158), n[+Irradiation] =131, p <0.0001).

### CBP regulates Tau acetylation upon DNA damage

To understand why Tau is hyper-acetylated upon DNA damage, we tested CBP levels. The acetylation of Tau is mainly carried out by p300 and CBP lysine acetyl transferases (KAT) (20,26). Our results show that CBP levels increase both in the nucleus and cytoplasm after irradiation (Fig. 4A and B). Accordingly, in SIRT6 deficient cells there are higher basal levels of CBP. Upon damage, both WT and S6KO cells reach their maximal level of CBP 3hr post-IR in the cytoplasm and 6hr in chromatin, and then decrease at 12h and 24h respectively (Fig. 3C-D). Curiously, endogenous levels of Tau-K174Ac increased 8 hrs post IR, like the effects on the CBP whose peak was at 6 hrs. Therefore, we hypostatized that inhibiting CBP could prevent Tau accumulation at chromatin. Indeed, the inhibition of CBP by C646 prevented nuclear accumulation of Tau in neuroblastoma cells in a dose dependent manner (Fig. 4E). To further understand whether CBP inhibition ameliorates Tau accumulation at chromatin under DNA damage, we used control and S6KO cells with irradiation in the presence of CBP inhibitor. Remarkably, Tau accumulation at chromatin was drastically reduced when the CBP inhibitor was administrated in both WT and S6KO cells, supporting our claim that the accumulation of Tau in the nucleus is driven by the enzymatic activity of CBP on DNA damage (Fig. 4F). Altogether, our results indicate that DNA damage and the lack of SIRT6 modulate CBP expression and location, which in turn impacts on the levels of Tau at chromatin.

**Figure 4.**
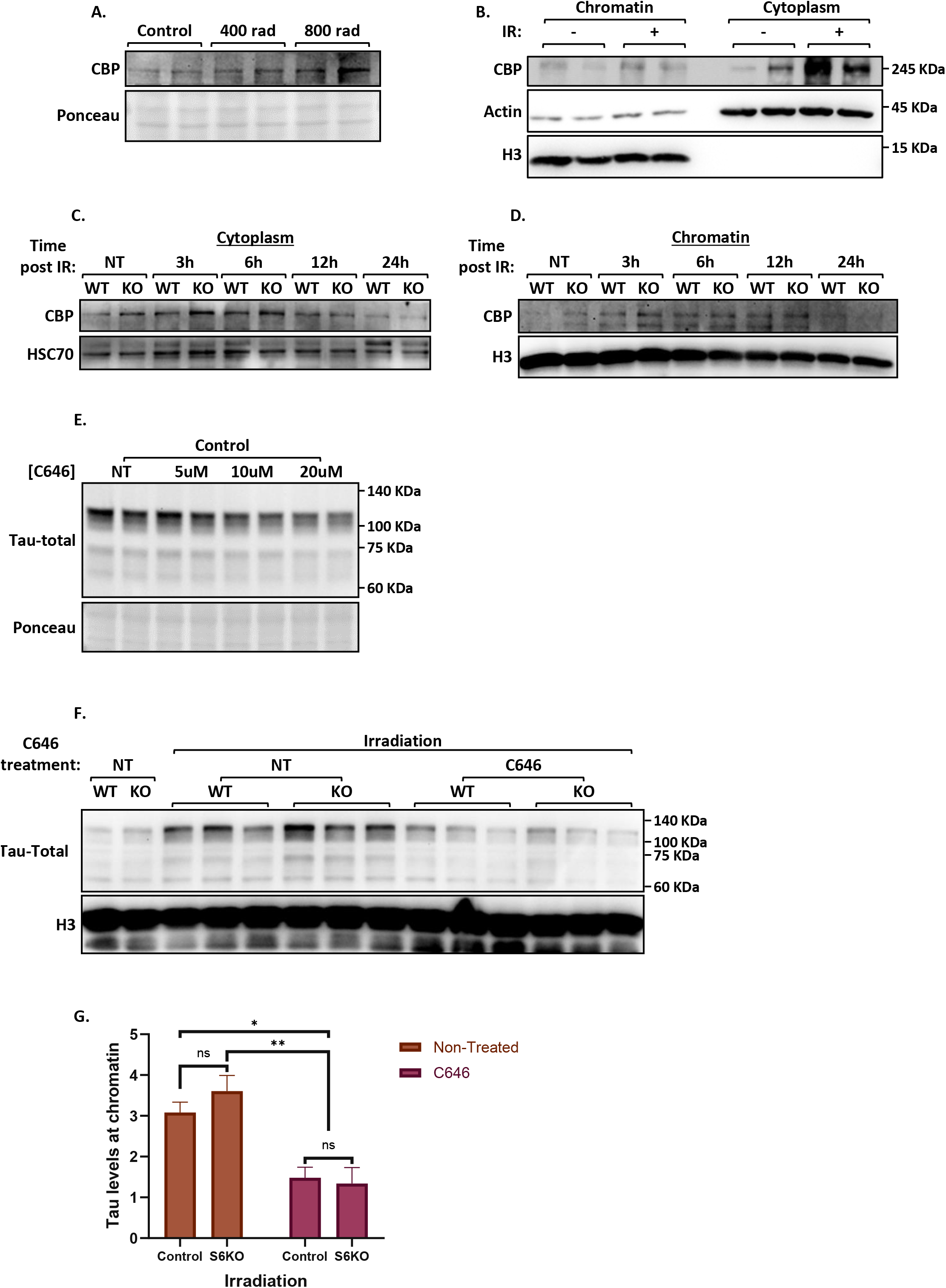
CBP expression is induced upon DNA damage. **(A)** Western blot of total extracts in SHSY-5Y cells subject to different irradiation doses. **(B)** Western blot of chromatin and cytoplasm fraction transfected with CBP and subjected to irradiation. **(C)** Time lapse of cytoplasmic and **(D)** chromatin extracts after irradiation **(E)** Western blot of Chromatin fraction of cells treated with increasing doses of C646 inhibitor **(F)** chromatin fraction of cells treated with C646 and irradiated with 4Gy **(G)** Densitometry of panel F. Error bars represent (SEM) (n[control +IR]= 3, n[S6KO+IR]=3, C646|n[control + IR]=3 and n[S6KO+IR]=3}), p<0.05, p<0.005)

### Tau acetyl mimic expression results in changes in ribosomal and translation gene expression

The increase in acetylated-nuclear Tau in pathological conditions, AD and SIRT6-KO cells/ mice, suggests that it has a pathological effect on the brain. To shed light on the physiological role of Tau-K174ac at the nucleus, we performed RNA-seq on neuron-like SHSY-5Y cells transfected with Tau-WT and acetyl mimic K174Q, in addition to the phospho-mimic S199E (regulated by SIRT6) and double mutant Tau-K174Q S199E. Our data show that several categories of protein translation were changed, as well as rRNA processing, the RNA biosynthetic process and ATP synthesis (Fig. 5A), suggesting global changes on translation capabilities and energy status of the cell. Comparisons between Tau PTMs and their interactions with each other (Acetyl mimic Tau-K174Q with Serine at residue 199 or a phosphomimic (Glutamate (E)) (Fig. 5B and SF5A) indicate that the main clusters were enriched for translation and ribosome biogenesis (see Fig. 4B and C, Cluster 1-3). Our data indicate that Tau-K174ac has an impact on cellular translation and energy production.

**Figure 5.**
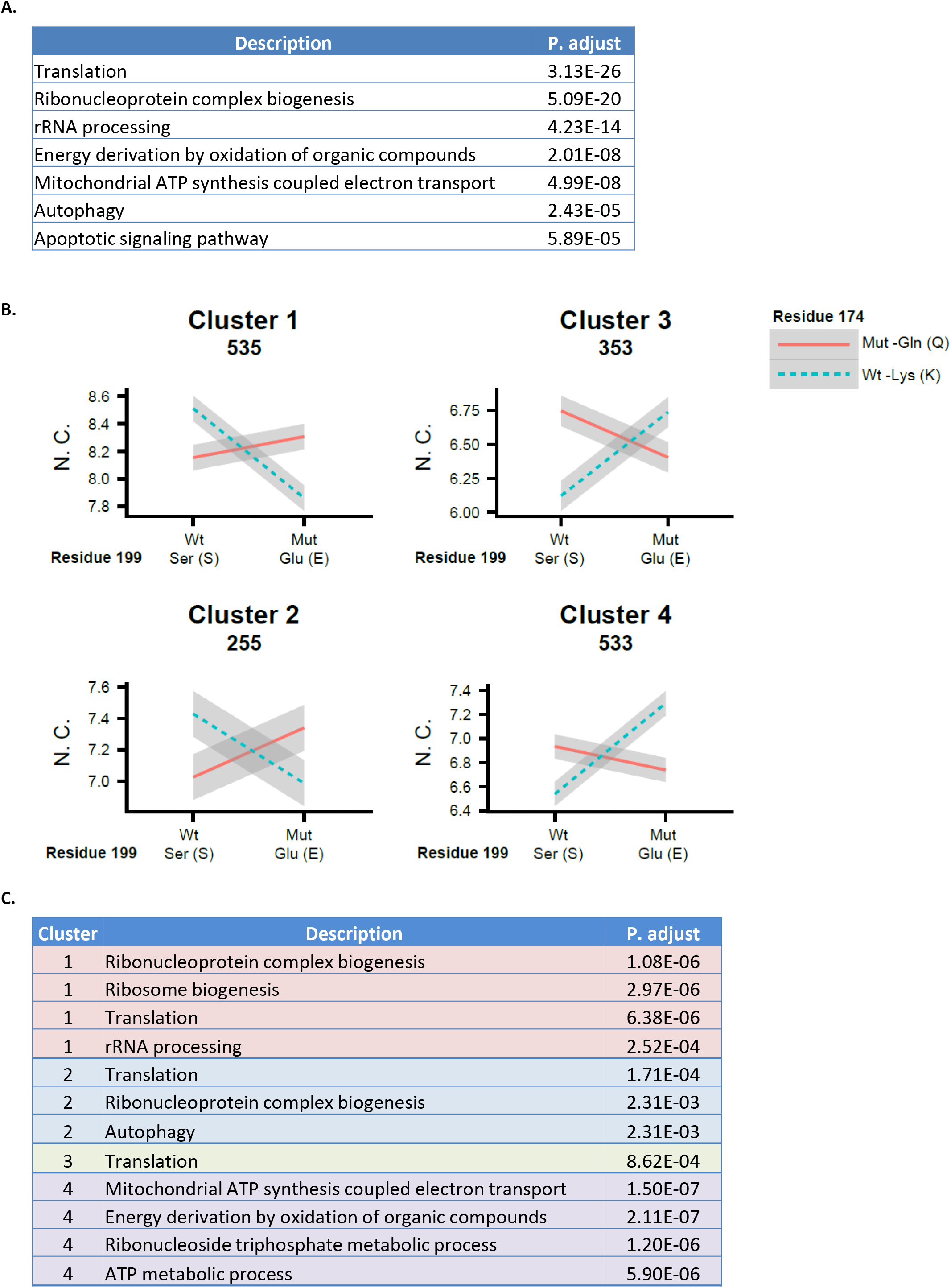
Tau 174Ac induces changes in ribosome biogenesis. **(A)** GO enrichment analysis of RNA-Seq from cells transfected with Tau-WT and Tau mutants, 199E, 174Q and 199E/174Q. **(B)** Clusters of Tau mutants, according to interaction between residue 174 and 199 (N.C. Normalized counts). **(C)** GO categories clustered by behavior.

### Tau-174Ac influences nucleolar activity and protein translation

RNA-seq results for Tau-174Ac suggest that translation should be affected at the transcription level as well. Notably, nucleolar function, size and number are directly correlated to protein synthesis; and an increase in these nucleolar parameters has a negative correlation with longevity. Interestingly, Tau was shown to be co-localized with the nucleolar protein Nucleolin (28). Moreover, Tau can bind AT-rich α-satellite DNA (28) and influence rRNA expression. Nucleolin is associated with active rDNA repeats, and nucleolar size correlates with the synthesis capacity of the cell to produce proteins (29). Thus, we measured nucleolar function by evaluating the size and number of the nucleolus, in addition to Nucleolin content (mean intensity). To test the effects of Tau-K174Q on the nucleolus in non-dividing and dividing cells, we used two systems: a primary neuronal culture and a SHSY-5Y neuronal-like cell line. Interestingly, in the primary neuronal culture Tau-K174Q increased Nucleolin content in the nucleolus without affecting its number or size (Fig. 6A and B, and SF6A-D). In accordance with these results, we found that the expression of Nucleolin increased in AD brains when compared to control (Fig. 6C). Moreover, SIRT6 has a positive correlation with Nucleolin expression in healthy brains, but this is lost in AD cases (Fig. S6F). An increment in Nucleolin has been shown to increase nucleolar activity through rDNA transcription. In line with these findings, Tau-K174Q also affected nucleolar function in neuroblastoma cells, however, unlike the primary brain culture, it increased the nucleolus number (Fig. 6D-E) instead of its Nucleolin content. Our results indicate that Tau-174Ac affects nucleolar activity increasing overall rRNA transcription, but different cells achieve this by distinct strategies. In the primary brain culture, it occurs via increased Nucleolin content in the nucleolus. In contrast, in neuron-like dividing cells, it occurs via increased nucleolar number (Fig. 6H).

**Figure 6.**
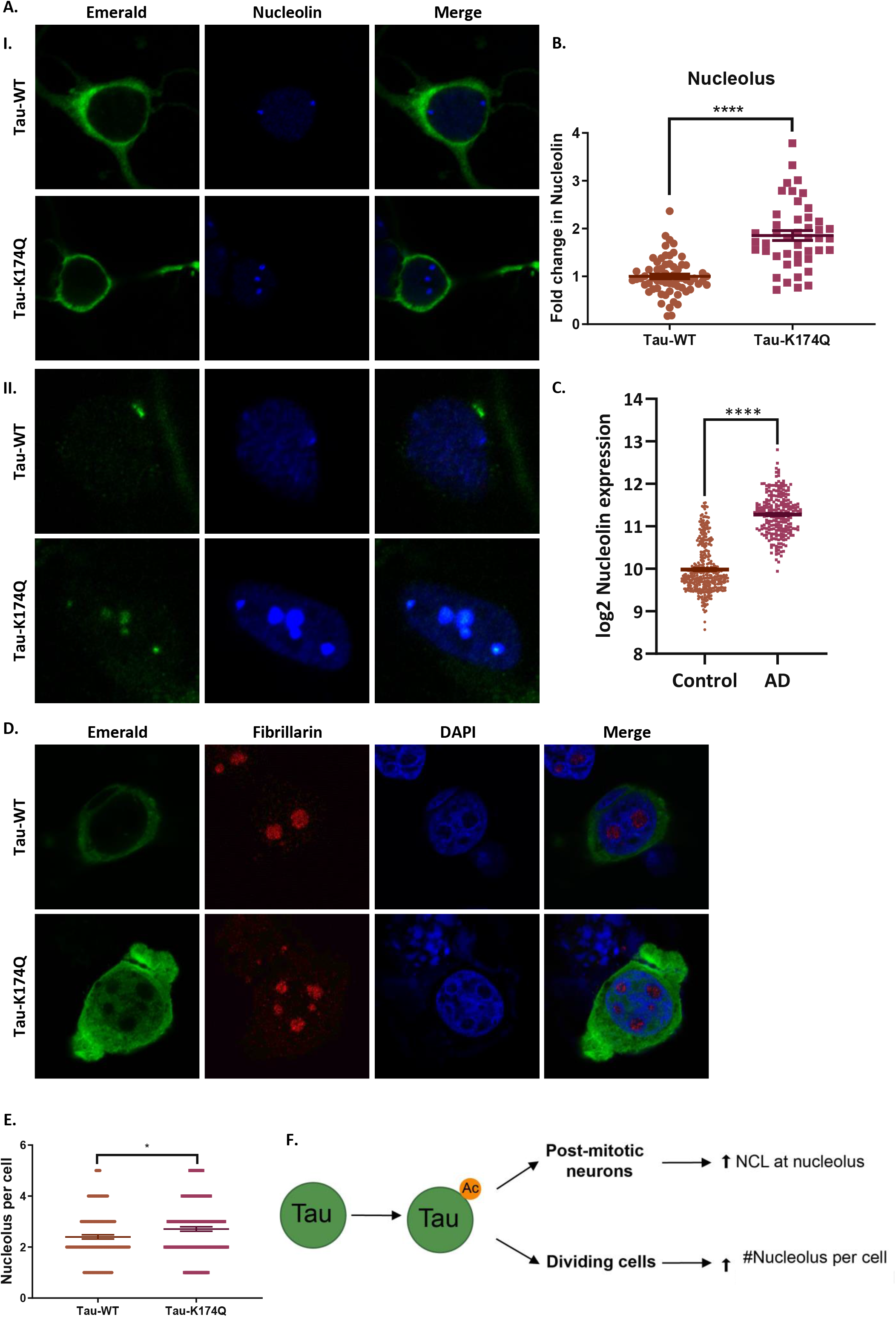
Tau 174ac induces nucleolar activation. **(A)** Immunofluorescence of neuronal primary culture infected with Tau WT and K174Q, lower and higher magnification (I and II respectively). **(B)** Fold change of nucleolin at nucleoli of neuronal primary culture Error bars are the Standard Error of the Mean (SEM), (n[Tau-WT]=63, n[Tau-174Q]=45, p<0.0001). **(C)** Nucleolin expresion analysis using R2: Genomics Analysis and visualization Platform (http://r2.amc.nl). Datasets: Alzheimer’s disease datasets: Brain-ADRC (Alzheimer’s Disease Research Center), Cotman 253, Alzheimer’s disease, and Salmon 74 for normal brains dataset (Core Transcript), Kang1340. **(D)** Immunofluorescence of Fibrillarin of neuroblastoma cells transfected with Tau WT and K174Q, **(E)** Nucleolus number per cell in five replicate experiments. Error bars are the Standard Error of the Mean (SEM), (n[Tau-WT]=158, n[Tau-174Q]=161, p<0.05). **(F)** Graphic representation of Tau-K174ac at nucleolus in post-mitotic neurons and dividing cells.

To confirm our hypothesis that rDNA is increased we measured RNA transcription in the nucleus and nucleolus in the presence of Tau-K174Q. For this purpose, we treated Tau transfected SHSY-5Y cells with 5-Fluorouridine (5-FU), which is incorporated in nascent RNA, and then measured rRNA synthesis by immunofluorescence (Fig. S7A) allowing us to localize nuclear versus nucleolar transcription. First, we look at 5-FU signal at the nucleus, where no differences between Tau-WT and Tau-K174Q were found at total RNA synthesis (Fig. S7B). Next, we used 5-FU mean intensity at nucleus to set a threshold for reference. We analyzed the sites with higher than average intensity which co-localized with the nucleoli (Fig. S7B-D). Confirming our hypothesis, rRNA transcription at nucleoli was increased in Tau-K174Q expressing cells when compared to Tau-WT (Fig. 7A-B). Moreover, active RNA synthesis sites were also increased in number and area (Fig. 7C-D). To test whether the increase in rRNA and nucleoli leads to an increase in protein synthesis, we performed a SUnSET assay where puromycin is incorporated to newly synthetized proteins and measured by immunoblot (30). Indeed, a significant increase in protein translation was detected when Tau-K174Q was expressed as compared to Tau-WT (Fig. 7E-F), suggesting that this acetylation is particularly important not only for its nuclear location, but also for the specific increase in rRNA synthesis.

**Figure 7.**
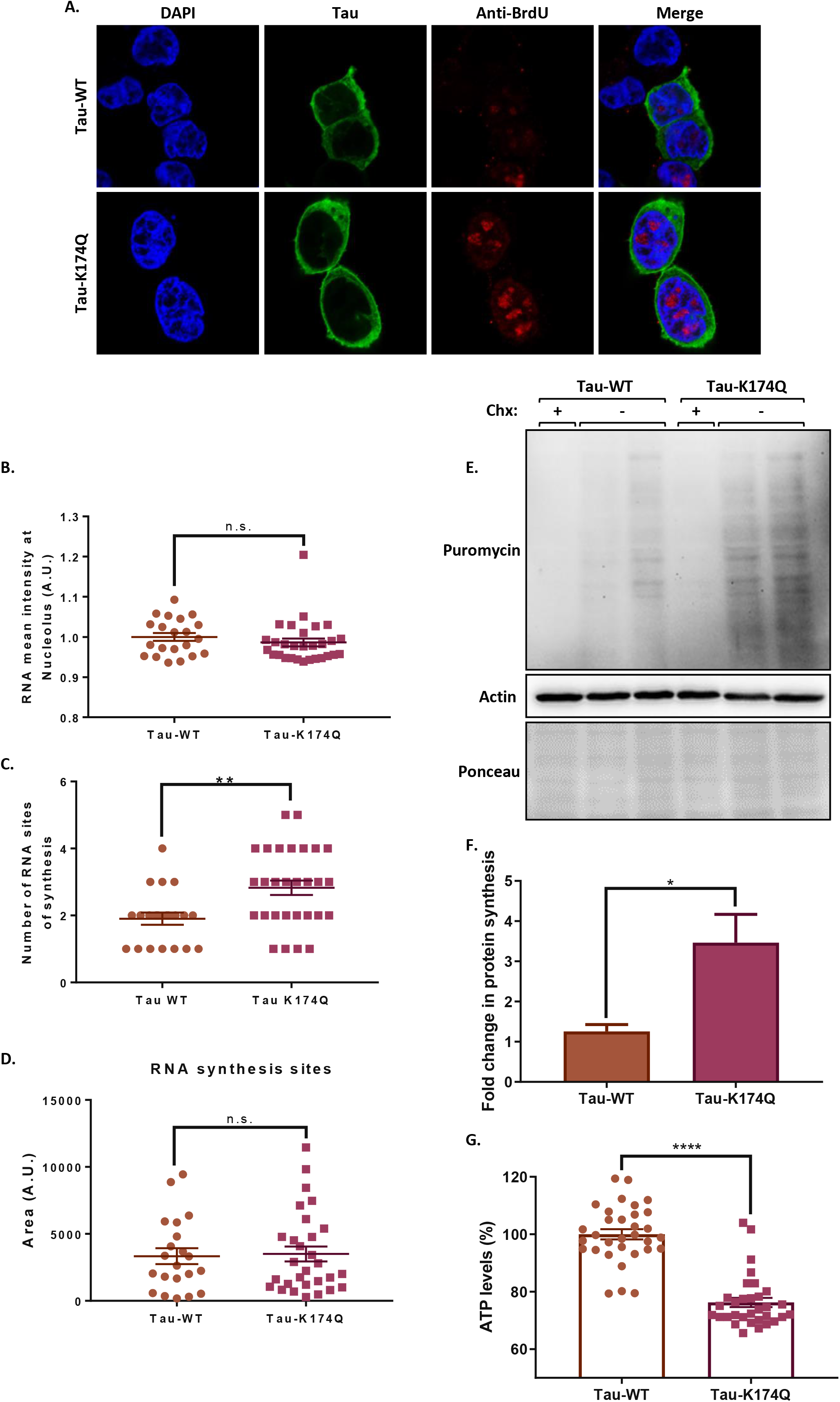
Tau 174ac impairs protein translation. **(A)** Immunofluorescence performed of SHSY-5Y cells transfected with Tau-WT and K174Q and treated with 5-FU. RNA synthesis was quantified using a nuclear threshold for 5-FU signal. **(B)** Mean intensity of 5-FU nucleoli per cell. **(C)** RNA synthesis sites per cell. **(D)** Combined area of RNA sites (passing the threshold) per cell. **(E)** SUnSET assay in SHSY-5Y cells transfected with Tau WT and K174Q. **(F)** Fold change in protein synthesis. Error bars represent (SEM) (n [Tau-WT] =4, n[Tau-K174Q] =4, p <0.05). **(G)** ATP levels in three independent experiments performed in SHSY-5Y cells transfected with Tau-WT and Tau-K174Q. Error bars represent (SEM) (n[Tau-WT] =32, n[Tau-K174Q] =32, p <0.0001). Cycloheximide (Chx).

Since protein translation in the cell is a high-energy consuming process, utilizing ~80% of the cell’s ATP, increased translation could affect cellular energy ultimately leading to its exhaustion (31). Therefore, we measured mitochondrial activity in cells expressing Tau-WT or Tau-K174Q. For that purpose, we used CellTiter-Glo^®^ to measure ATP levels and the results were normalized by total protein amounts (as an equivalent for cell number). We observed a 23.75% reduction of ATP when Tau-K174Q was expressed as compared to Tau-WT, strongly suggesting that an increase in protein translation indeed depletes the energy levels of cells (Fig. 7G). In agreement with this, we also observed AMPK activation (hyper-phosphorylation at 172) in cells transfected with Tau-K174Q (Supp. Fig. 7E). AMPK its activated in response to the increase in AMP:ATP and ADP:ATP ratios which occurs under energy stress (32). Taken together, these results indicate that Tau-K174ac induces rDNA relaxation, nucleolar expansion, and an increase in translation capacity, ultimately leading to the energy exhaustion of the cells.

Overall, we have shown that Tau is acetylated at residue K174 upon genotoxic stress leading to its nuclear localization by CBP. This leads to global changes in gene expression, mainly in genes related to energy usage and translation, allowing the cell to cope with the challenges imposed by DNA damage. However, when Tau deacetylation is prevented, either by locking off the residue 174 as in the case of Tau-K174Q or by the depletion of SIRT6, Tau accumulates in the nucleus leading to translational changes and increases nucleolus capacity by allowing an increment in rRNA synthesis and protein translation which eventually exhausts ATP levels (Fig. 8). It is important to highlight that these changes are also observed in a model of accelerated ageing and neurodegeneration such as brSIRT6KO mice and in brains of AD patients.

**Figure 8.**
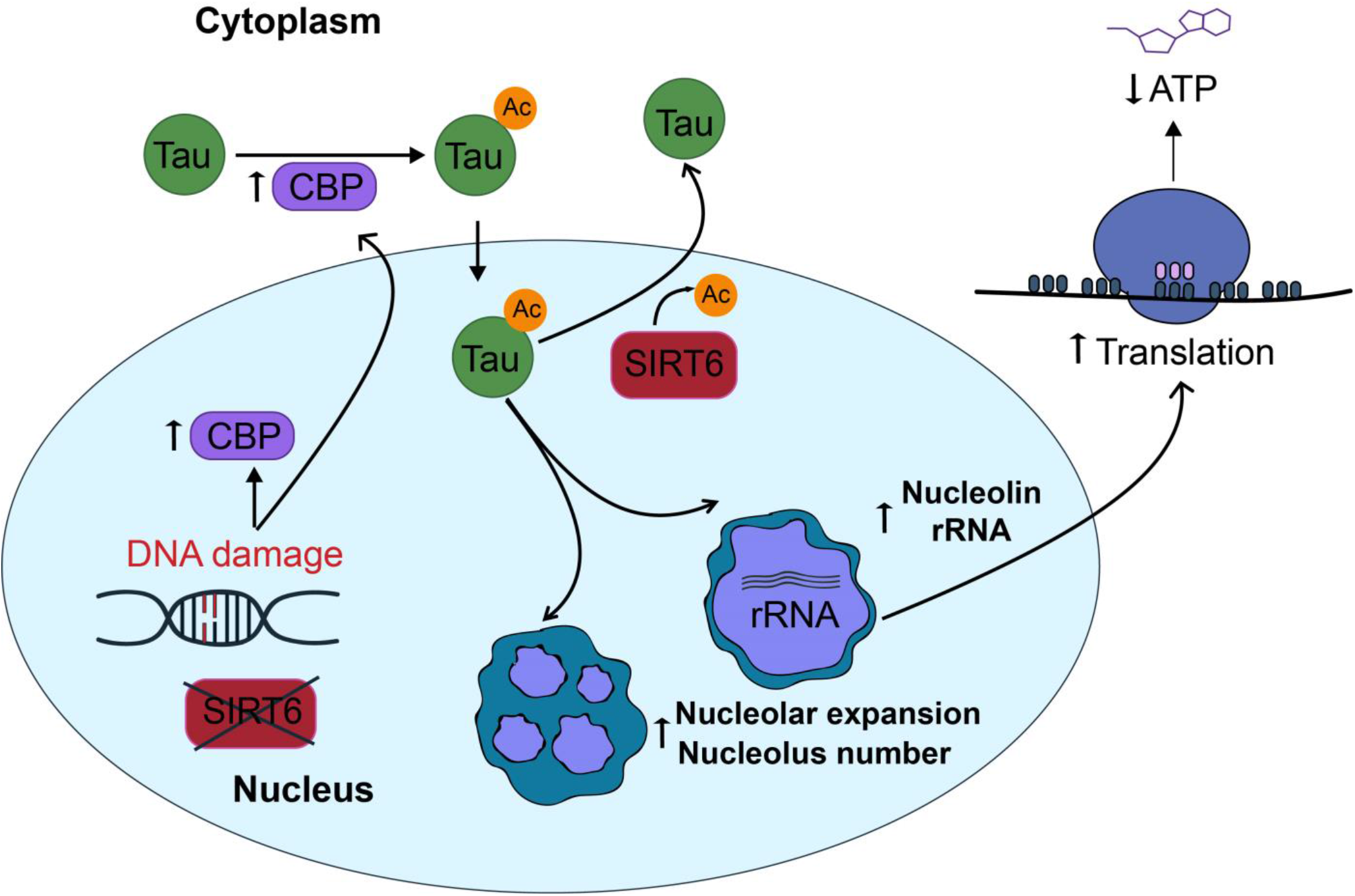
Proposed mechanism. During DNA damage CBP is upregulated in the cells, both at nucleus and cytoplasm. CBP acetylates Tau at Lys 174 in the cytoplasm, facilitating Tau 174ac translocation to the nucleus. In the nucleus Tau174Ac increases nucleolar activity leading to increase transcription of rRNAs and other mRNAs important for translation, RNA biogenesis and energy production. In the nucleus SIRT6 negatively regulates Tau through deacetylation at 174ac, allowing Tau to exit the nucleus. However, during ageing, or constant DNA damage, SIRT6 levels decrease, Tau and CBP levels increase leading to an increase in Tau 174Ac. Tau174ac accumulates in the nucleus-nucleolus, resulting in changes in gene expression, increase protein production and ATP reduction.

## Discussion

The accumulation of Tau in the cytoplasm and its role in neuropathology have been extensively studied. However, the various roles of Tau in the nucleus are just beginning to emerge as critical in AD. Most of the focus has been given to mutated Tau, even though most cases of neurodegeneration are sporadic (95%) and the changes are probably linked to cellular signaling and ageing (6). In this work, we show that under stress conditions, like DNA damage or models of accelerated ageing such as SI RT6-deficient brains, the nuclear localization of Tau is increased. This is in part due to the acetylation at residue K174, which directs Tau towards the nucleus. Once there, Tau-K174ac changes the transcription profile of the cells increasing their translation capacity and leading to energy depletion.

### SIRT6 regulates Tau stability and nuclear location through the deacetylation of residue 174

We previously showed that SIRT6 regulates Tau levels through its phosphorylation by GSK3 activation (33). We found that SIRT6 also regulates nuclear Tau through deacetylation. Nuclear Tau levels were increased in the brain tissue of brS6KO, old mice, and Alzheimer’s disease patients, supporting the role of nuclear Tau as a pathological driver and SIRT6 as its modulator. It is intriguing that the immunohistochemistry of Tau-K174ac in AD patients showed that it is mainly localized in the nucleus (Fig. 2–3 and SF2–3). Our data suggest that Tau acetylation at Lys174 allows the shuttling of Tau to the nucleus, where SIRT6 restricts its permanence through its deacetylase activity (Fig. 1 and 2). It is worth noticing that SIRT1 also targets Tau-174Kac ameliorating Tau spreading in Tau-PS301 transgenic mice (34). Tau-K174ac deacetylation by SIRT1 and 6 raises the possibility that Tau can be differentially regulated in the cytoplasmic and nuclear compartments by different Sirtuins under different cellular contexts (35). In line with this hypothesis, Hou et al. found that NAD^+^ supplementation ameliorates AD phenotype in 3xTGAD/Polβ^+/-^ in a SIRT6 dependent manner (36). Remarkably, the absence of either SIRT6 or Tau-K174Q mutant lead to fragmentation and accumulation of Tau in the nucleus, suggesting that a failure in K174 deacetylation can trigger Tau self-cleavage activity and ulterior toxicity (Fig. 2) (37,38). Tau fragmentation could be an outcome when the cell attempts to ameliorate Tau’s effect on the nucleus thus generating toxic fragments.

Others have shown that the amount of Tau correlates to those of H3K9ac in the dorsolateral prefrontal cortex of human samples (39,40). Moreover, they showed that Tau is associated with H3K9ac in open chromatin, suggesting that both Tau-K174Ac and H3K9ac allow chromatin opening. Interestingly, this modification is one of the main targets of SIRT6. In fact, when SIRT6 is not present, H3K9ac and Tau-K174ac accumulate, resulting in an overly relaxed chromatin that could alter gene expression. This relaxation may lead, for example, to transposon activation, as can be seen both in SIRT6 deletion and Tau overexpression, which in turn increase genomic instability (6).

### Tau-174Ac is translocated into the nucleus upon DNA damage

SIRT6 is critical to ensure DNA repair (10,41). We have recently shown that it acts as a double strand breaks (DSB) sensor (42) and prevents genomic instability. We have previously established a connection between DSB signaling, Tau stability and its accumulation through GSK3 activation and Tau phosphorylation (23,26). Since DNA damage accumulation could cause neurodegeneration in aged population, we tested whether this signal could increase nuclear Tau and acetylated Tau at residue K174. Our results showed that Tau-K174ac is induced upon irradiation, increasing its presence in chromatin over time (Fig. 3). Given their impaired capability to repair DNA and deacetylate Tau-K174ac, S6KO cells show higher accumulation of Tau in the nuclear compartment, further supporting our claim that acetylation is a driver of Tau nuclear accumulation (Fig. 3). Under normal conditions, Tau shuttles between cytoplasm and nucleus, where it has roles in DNA protection and chromatin relaxation (43). However, under constant stress such as DNA damage accumulation — which occurs as we age (44,45) — and under reduced SIRT6 levels (again seen in old age or AD patients), Tau becomes hyper-acetylated and remains in the nucleus. Overall, our results support the protective role of SIRT6 in neurodegeneration and its important driving role in DNA damage in neurodegenerative Tauopathies.

### CBP regulates Tau upon DNA damage

Tau acetylation regulates important aspects of Tau pathophysiology such as aggregation, spreading, microtubules stabilization and its location (29). Intriguingly, the acetylation of Tau is carried out by the nuclear proteins CBP and p300. How CBP/p300 meets cytoplasmic Tau for its acetylation and the context in which this occurs remains unaddressed. Our results showed that either DNA damage or SIRT6 deletion increased CBP levels in the nucleus and cytoplasm (Fig. 4 and SF4). SIRT6-KO cells are deficient in their ability to repair DNA and CBP is a key player on DSB repair (46,47), thus its overexpression might be the outcome of persistent DNA damage signaling and an important component of the cellular efforts to mitigate DNA damage. Our results show that preventing CBP activity can also ameliorate Tau accumulation and nuclear localization. Besides, our data suggest that CBP may be the link that connects DNA damage with Tau shuttling under DSB (Fig. 4 and SF4).

### Tau-174Ac influences nucleolar function

In neurons, Tau can be found in the nucleolus where it may play a role on heterochromatinization of rDNA, rRNA transcriptional regulation, maturation and processing (47). In the nucleolus, Tau protein interacts with TIP5 repressing rDNA transcription. Our results show that Tau acetylation at Lys-174 induces nucleolar activity in dividing and non-dividing cells. This was measured by an increase in the number of nucleolus, and increased Nucleolin amounts in the nucleolus respectively. Nucleolar size increase has been shown to negatively correlate with lifespan in models of premature ageing and progeroid-like syndromes (47). Interestingly, one of the known functions of Nucleolin is to decrease TIP5 and HDAC1 occupancy at rDNA. These proteins are components of the NoRC silencing complex, therefore, an increase in Nucleolin can lead to rDNA de-repression (48).

Nuclear Tau is highly sensitive to cellular stressors (48), thus different stressors may modulate Tau’s PTMs. Tau could have achieved various responses depending on the specific modification. For example, DNA damage will induce Tau acetylation at residue K174, affecting nucleolar regulation. In contrast, Maina et al (48) reported that nuclear Tau is reduced when cells are treated with Aβ42 oligomers, correlating it with non-phosphorylated Tau. These changes reduced TIP5, FBL, rRNA production and protein translation in their model (48). In contrast, our results suggest that Tau-K174ac positively regulates rRNA production and protein synthesis, highlighting the importance of nuclear Tau and the differential roles of its PTMs. Altogether, our results suggest that Tau acetylation can regulate its nuclear/ nucleolar functions, affecting chromatin in the nucleolus.

Moreover, our results show that Nucleolin expression is highly increased in AD patients, while SIRT6 levels dramatically drop (Fig. S6E). It is interesting to mention that SIRT6 and Nucleolin seem to be co-expressed in healthy human brain samples, however, this correlation is lost in AD patients (Fig. S6F).

### Tau 174 induces changes in protein synthesis

The comparison between the changes in gene expression caused by Tau-WT or Tau-K174Q resulted in different clusters of behavior, denoting that a simple PTM can affect the roles of Tau. Tau-WT is less stable than Tau-K174Q, therefore, some changes could be due to an increase in the protein amount. However, the cluster analysis showed a clear difference in the pattern of expression and not only an incremental change in the same categories. Protein interactions are commonly regulated by posttranslational modifications hence it is possible that Tau-acetylation influences the nuclear proteins it interacts with, affecting gene expression as an outcome. Our results showed changes in genes related to RNA and protein synthesis, energy consumption, and autophagy (Fig. 5 and SF5). Accordingly, protein translation was increased in cells expressing the acetyl mimic, Tau-K174Q (Fig. 7). Tau has been reported to modulate translation before. However, those studies refer to Tau as a cytoplasmic regulator of protein synthesis, which is achieved through its direct binding to the ribosomes, thus decreasing translation (48). Our findings show that Tau-K174ac controls translation through gene expression. Changes in gene expression are essential to allow the cell to cope with the DNA damage stress. Nevertheless, when the damage is sustained or SIRT6 is depleted, it will lead to an exacerbated response encompassed by protein overload. An increase in protein synthesis without an accompanied increase in chaperons may contribute to protein aggregation as seen in AD patients. Interestingly, we also observed through our RNA-seq an increase in the activation of various pathways from the “Unfolded Protein Response” when Tau174Q is expressed, as seen in various neurodegenerative diseases (Fig. S5B). Moreover, protein translation is necessary for learning and memory formation (48), thus the misregulation of such process could lead to cognitive deficiencies. Consequently, changes directed by increased Tau-K174ac would alter the global pattern of protein synthesis and protein misfolding, and together would lead to an increase in energy consumption, which may result in a neurodegenerative phenotype.

Several age-related neurodegenerations have in common changes in Tau levels and PTMs^35^. Much is known of Tau as a toxic aggregate and a bystander that becomes toxic due to PTMs, aggregates or soluble fractions. Here, we show that an increase in Tau-K174ac provokes toxicity when it accumulates in the nucleus; not as a bystander but as a protein with important roles in chromatin protection and the cope with DNA damage. However, prolonged stress can result in its nuclear accumulation, impeding the turning off of its nuclear function. Given that Tau can be modified at several residues and, probably, in different combinations, it is possible that there are various “Tau PTMs-barcodes” that differ among Tauopathies, resulting in a myriad of phenotypes associated with them.

In ageing, and even more in AD patients, DNA damage accumulates in the brain. DNA damage signaling becomes constant, leading to Tau-K174ac accumulation, translocation and deleterious nucleolar expansion. Our findings show for the first time a PTM that results in Tau translocation into the nucleus, and reveal its function in rRNA expression which increases protein synthesis and ATP exhaustion. Paradoxically, these brains have lessened DNA repair capacity, due to the lack of SIRT6, therefore, they have increased DNA damage signaling and CBP expression, which in turn leads to an increase in Tau-K174ac. Tau is then constantly acetylated and translocated into the nucleus where it gets trapped, again by the lack of SIRT6, which is also required to remove Tau from there. This gives place to a vicious cycle of accumulation of nuclear Tau, ultimately resulting in neurodegeneration.

## Acknowledgements

This work was supported by The David and Inez Myers foundation, ISF 188/17 and by the High-tech, Bio-tech and Negev fellowships of Kreitman School of Advanced Research of Ben Gurion University.

---

**Supplementary Figure 1.**
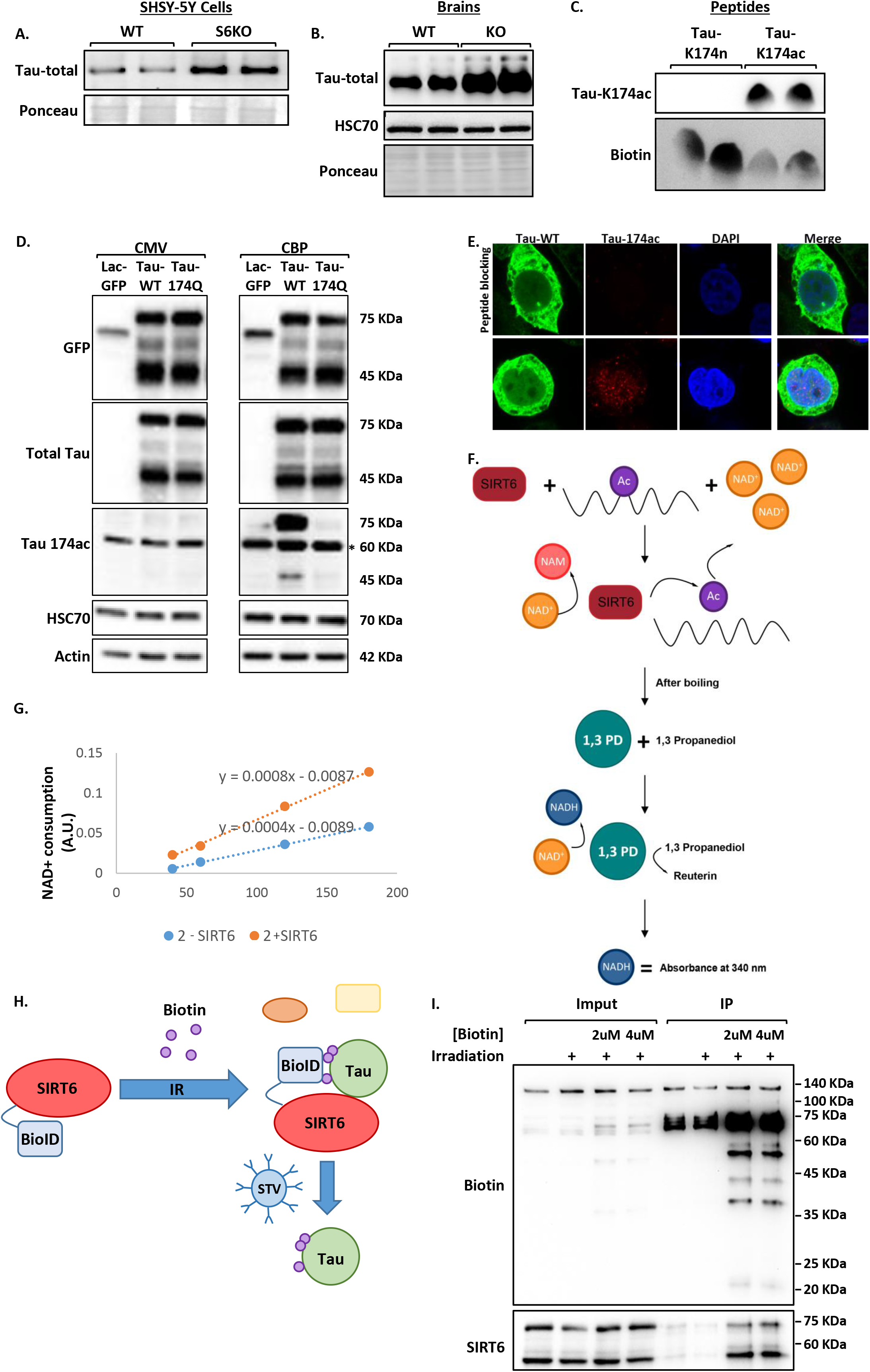
**(A)** Western blot of SHSY-5Y cells WT and S6KO cells showing Tau levels. **(B)** Western blot of mice brains WT or KO for SIRT6 (Nestin-Cre). **(C-E)** Validation of Tau-174ac antibody specificity: **(C)** Western blot of control and Tau-174ac peptides using Tau-174ac antibody. **(D)** Western blot of cells co-transfected with Tau-K174Q and CBP. **(E)** Immunofluorescence of Tau-174ac in cells transfected with Tau-WT. Cells were blocked with Tau-174ac peptide as a specificity control. **(F)** Schematic representation of NAD+ consumption assay. **(G)** NAD+ consumption assay using H3K56ac peptides showing rates of conversion of NAD to NADH represented as (1-NADH). **(H)** Graphic representation of SIRT6-BioID proximity assay: Proreins in close proximity are biotinylated by: himera SIRT6-BioID, this was used to detect transient SIRT6 interactors. **(I)** Immunoprecipitation of biotinylated proteins performed on cells transfected with SIRT6-BioID and irradiated with 4Gy.

**Supplementary Figure 2.**
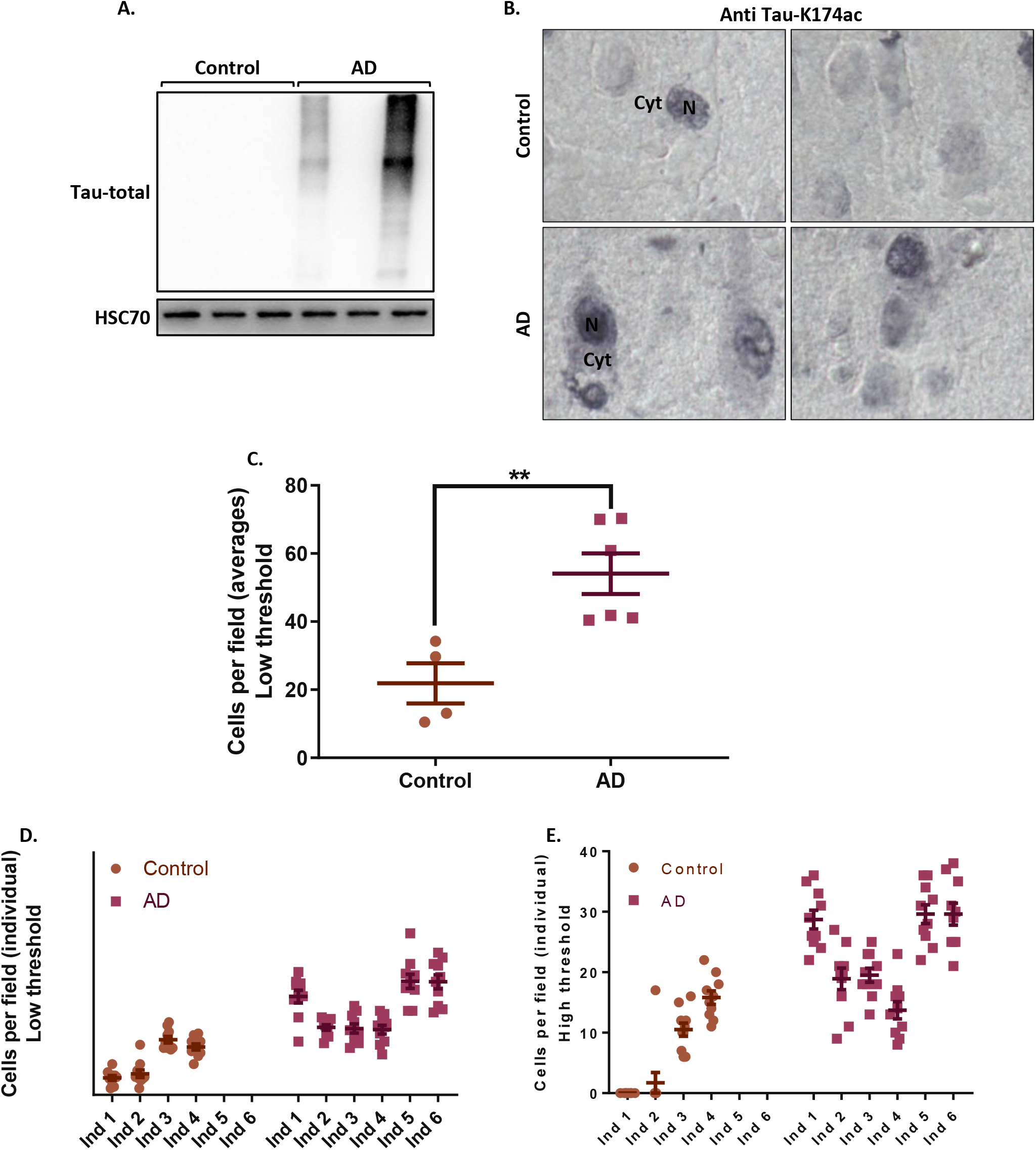
**(A)** Western blot of cytoplasmic fraction from samples of brains from Alzheimer disease patients or non-demented controls. **(B)** Immunohistochemistry of Control and AD brains (N: Nucleus; cyt: Cytoplasm). **(C-D)** Manual quantification of Tau-174ac positive cells per field (n[Control]=4, n[AD]=6). **(C)** Average of Tau-174ac positive cells per field, non-stringent threshold. **(D-E)** Tau-174ac positive cells per field, using non stringent **(D)**/stringent **(E)** threshold Error Bars are the Standard Error of the Mean (SEM) (n[Control]=4, n[AD]=6, 10 images were quantified per individual and their values graphed at D and E, p <0.001).

**Supplementary Figure 3.**
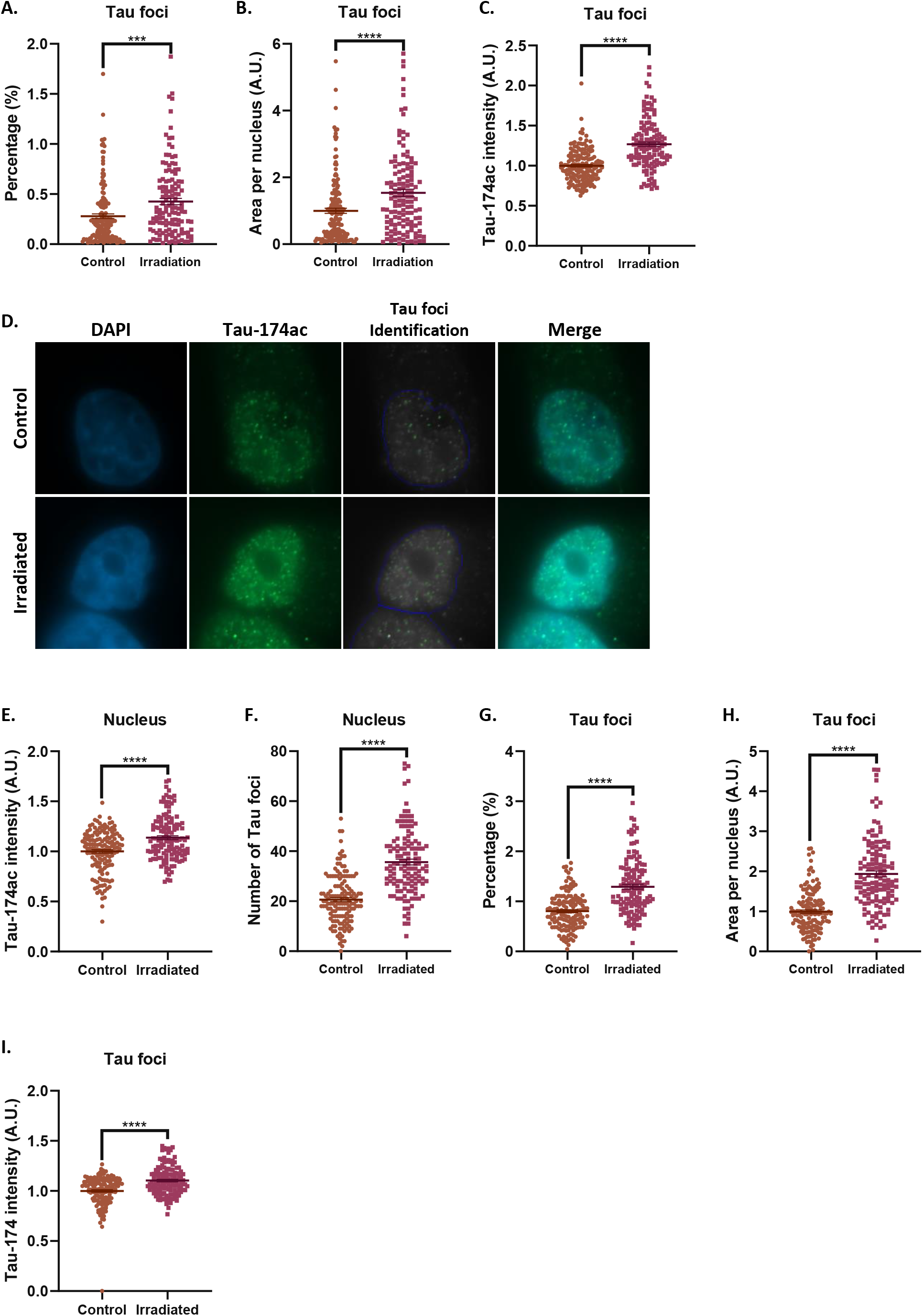
**(A-C)** Tau-174ac foci quantifications in primary neuronal culture with or without irradiation **(A)** percentage of Tau foci/nucleus area. **(B)** Tau are of the foci per nucleus. **(C)** Tau-174ac mean intensity at foci. Error bars are the Standard Error of the Mean (SEM) (n[-Irradiation] =158), n[+Irradiation] =131, p <0.0001). **(D-I)** Immunofluorescence and quantifications of Tau-174ac in SHSY-5Y cells (quantification of two independent experiments). **(D)** Immunofluorescence of Tau-174ac performed in SHSY-5Y cell line -/+ irradiation. **(E)** Tau-174ac mean intensity at nucleus. **(F)** Number of Tau foci per nucleus. **(G)** percentage of Tau foci/nucleus area. **(H)** Area of Tau foci per nucleus. **(I)** Tau-174ac mean intensity at foci. Error bars represent (SEM) (n[control]=145), n[+Irradiation] =129, p <0.0001).

**Supplementary Figure 4.**
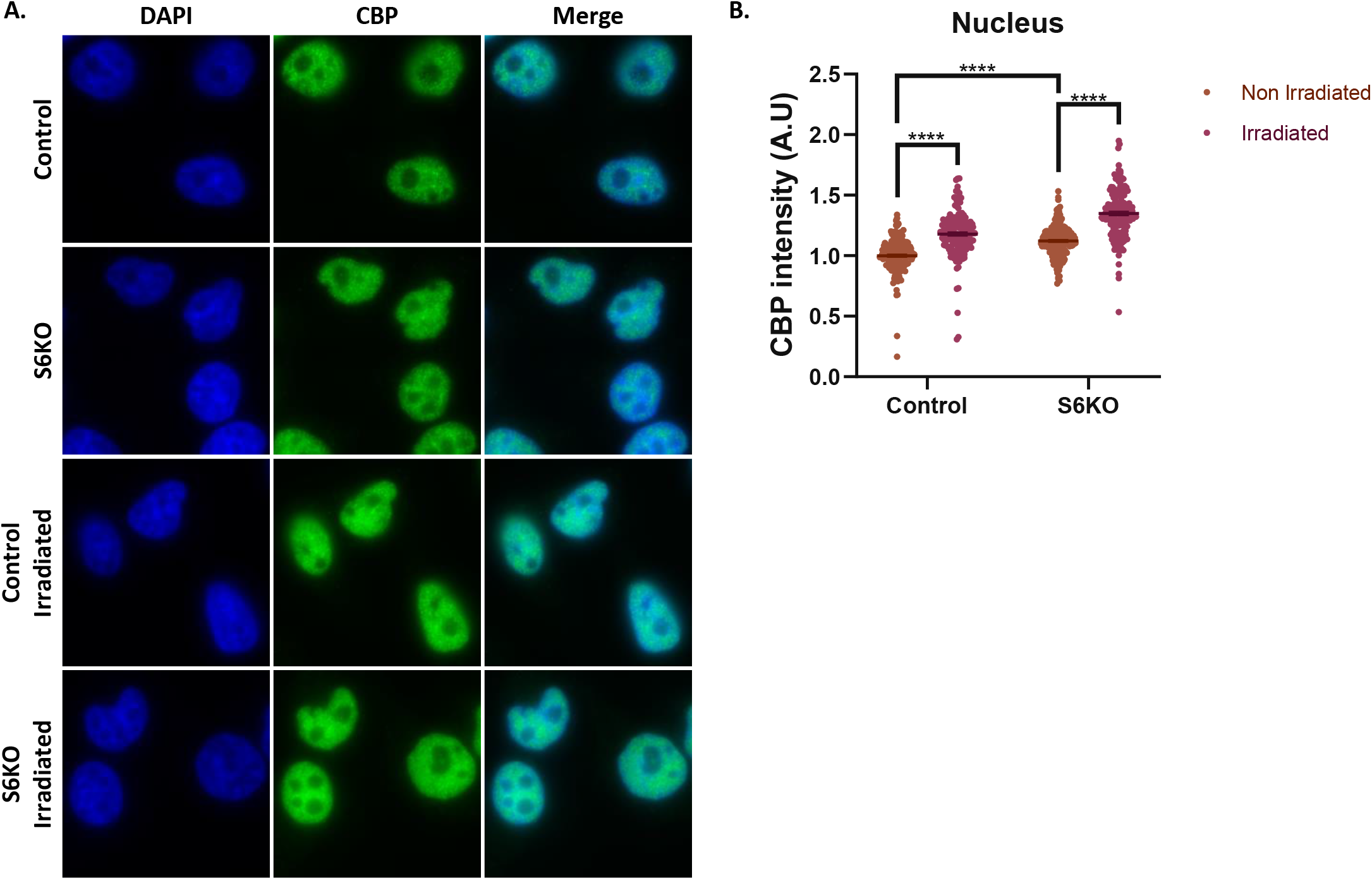
**(A)** CBP immunofluorescence in WT and S6KO SHSY-5Y cell line -/+ irradiation. **(B)** CBP mean intensity at nucleus. Error bars represent (SEM) (n[control]=167, n[S6KO] =165, n [control +IR] =152 and n[S6KO+IR] =164, p<0.0001).

**Supplementary Figure 5.**
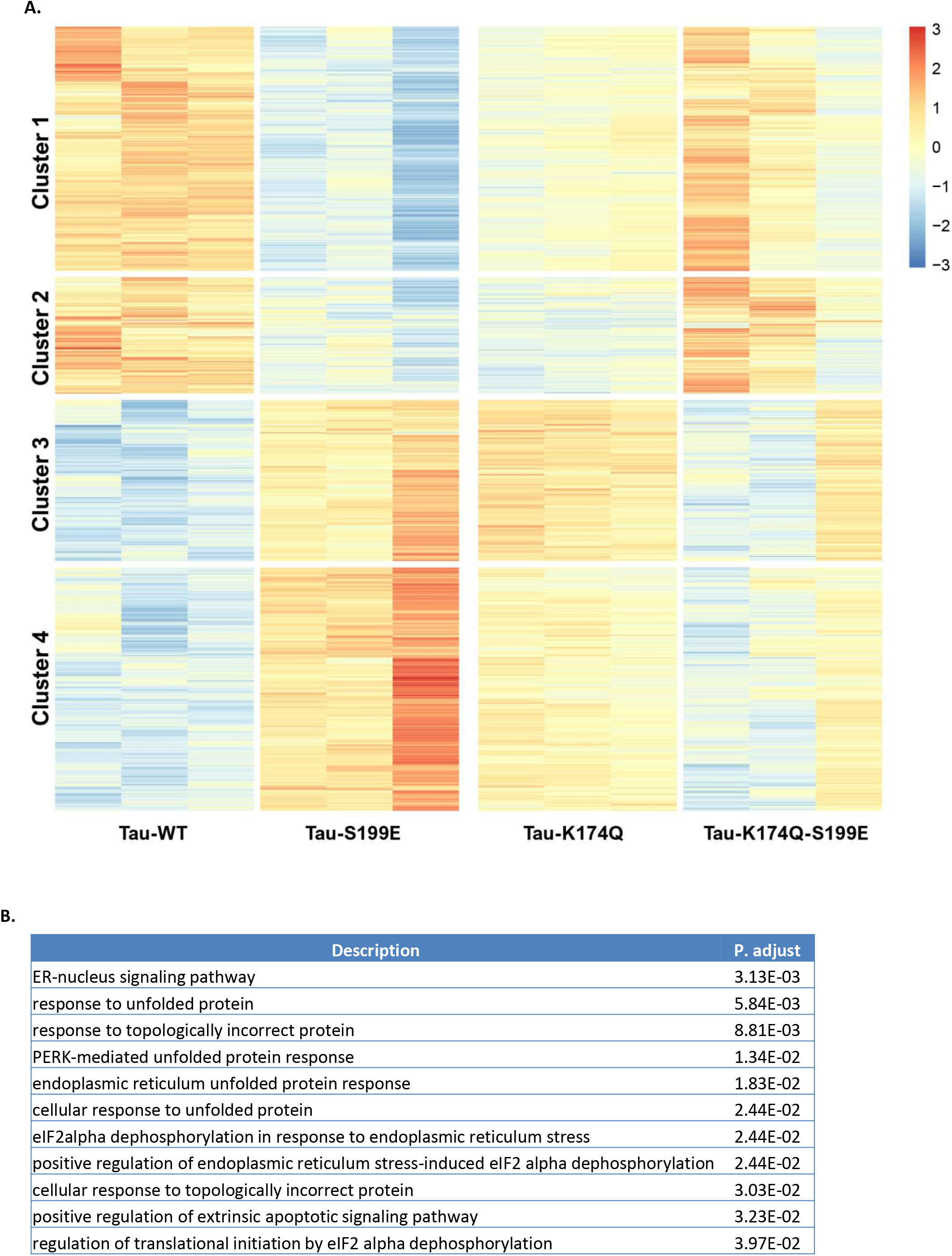
**(A)** RNA-seq clustering heatmap of SHSY-5Y cells transfected with Tau-WT and Tau mutants. **(B)** GO enrichment analysis highlighting categories related to protein misfolding in cells expressing Tau-WT and K174Q mutant.

**Supplementary Figure 6.**
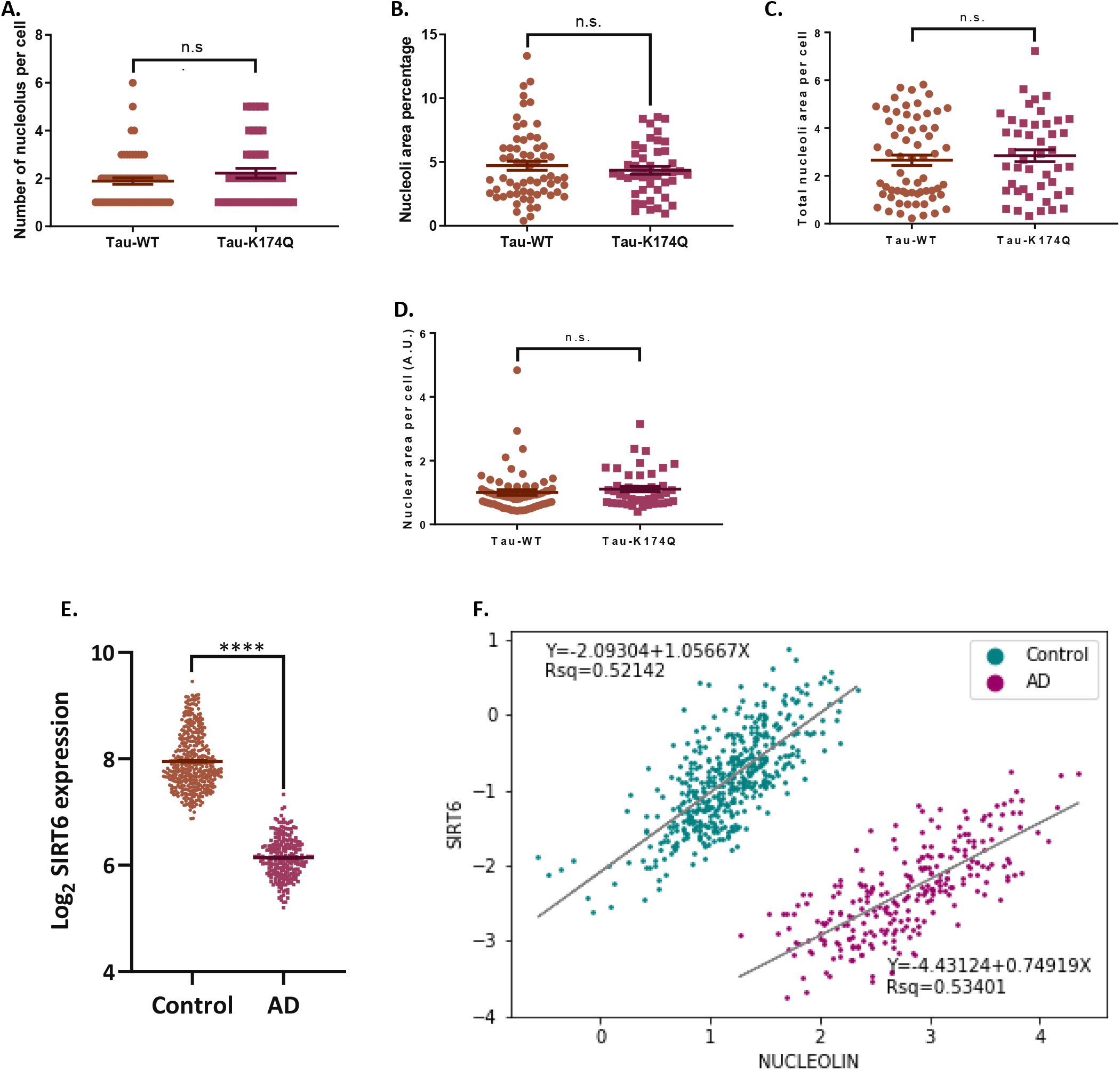
**(A-D)** Manual Quantifications of neuronal primary culture infected with Tau-WT and-K174Q (A) Number of nucleolus per cell **(B)** Percentage of nucleoli area normalized to its nucleus **(C)** Total nucleoli area per cell **(D)** Nuclear area per cell. **(E)** SIRT6 expression analysis using R2: Genomics Analysis and visualization Platform (http://r2.amc.nl). Datasets: Alzheimer’s disease datasets: Brain-ADRC (Alzheimer’s Disease Research Center), Cotman 253, Alzheimer’s disease, and Salmon 74 for normal brains dataset (Core Transcript), Kang1340. Error bars represent (SEM), (n[Control]=404, n[AD]=253, p<0.0001). **(F)** Comparison of SIRT6 and nucleolin expression using R2: Genomics Analysis and visualization Platform (http://r2.amc.nl). Datasets: Alzheimer’s disease datasets: Brain-ADRC (Alzheimer’s Disease Research Center), Cotman 253, Alzheimer’s disease, and Salmon 74 for normal brains dataset (Core Transcript), Kang1340. Samples were filtered by age (age>20 years). Normalization was achived using Python Software foundation (version 3.7). ANOVA test was done on H4 expression, extracted from the datasets, being this not significant (p>0.05). Normalization was achieved by subtracting the log2 of housekeeping genes to each sample (log2[gene])-log2[H4]= log2[gene/H4]). Afterwards, linear regression was done on each group separatly and were compared using Real Statistics Resource Pack Software (release 6.8). Copyrigth (2013-2020) Charles Zaiontz. http://www.realstatistics.com/free-download/real-statistics-resource-pack/. Data was plotted using python. Arbitrary Units (A.U).

**Supplementary Figure 7.**
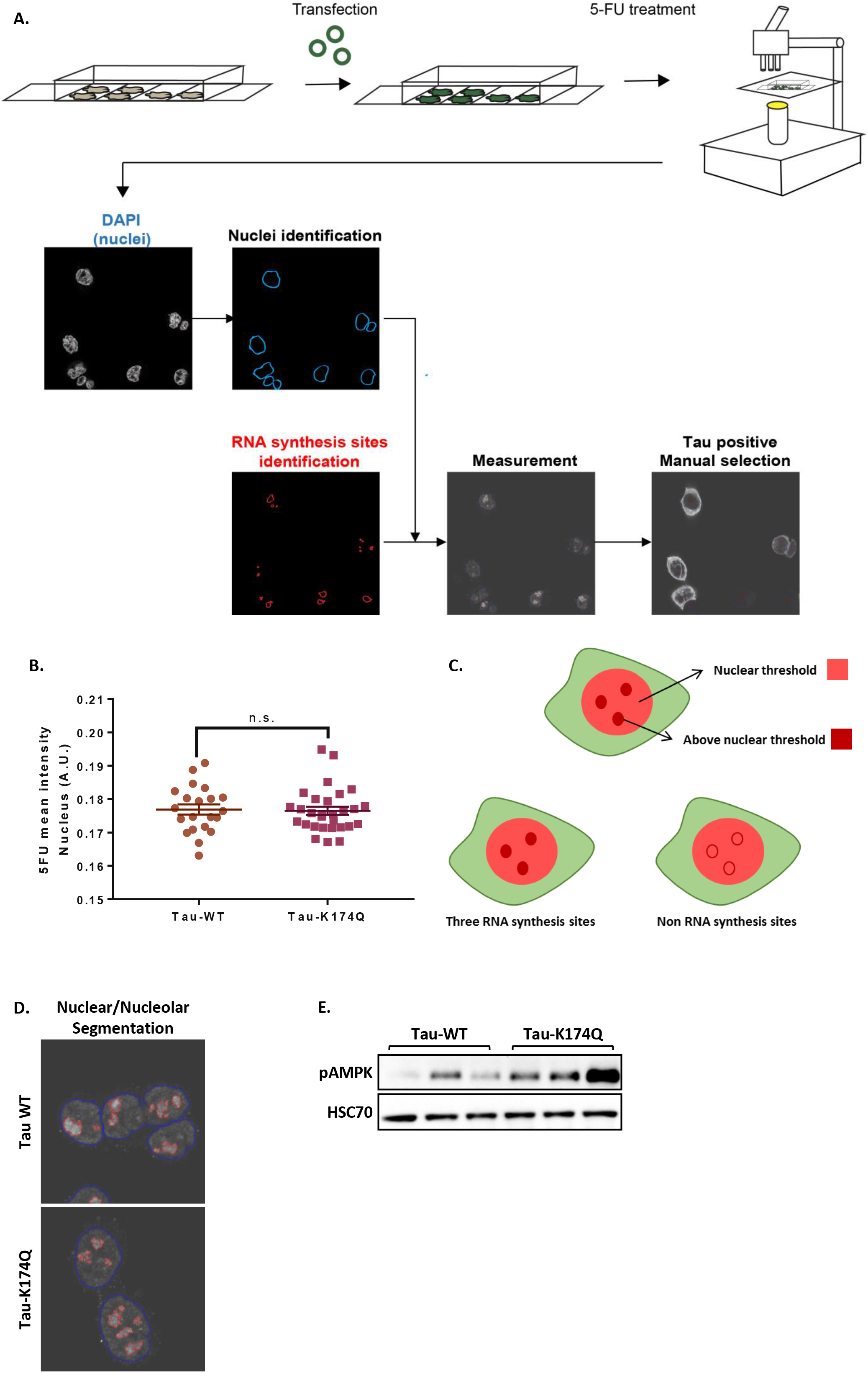
**(A)** Schematic representation of 5FU experiment and analysis. Cell were transfected with Tau-WT and K174Q mutant. One day pos-transfection, cells were treated with 5-FU for 30 minutes. Pictures were analyzed with the Cell Profiler Software by first recognizing the nuclei in each cell by DAPI staining, then we analyzed the far-red channel (Alexa Fluor 647 for anti-BrdU), to identify 5FU marked cells and measured nucleolus number and intensity. Next the software associates 5-FU/RNA sites to each nucleus and acquire its measurements. Finally, data was manually cleaned by selecting only Tau positive cells. **(B)** Mean intensity of 5-FU at nucleus **(C)** Scheme of criteria for the recognition for 5FU/RNA sites. **(D)** Nuclear and 5-FU segmentation. **(E)** Western blot of pAMPK in cells transfected with Tau-WT and K174Q.

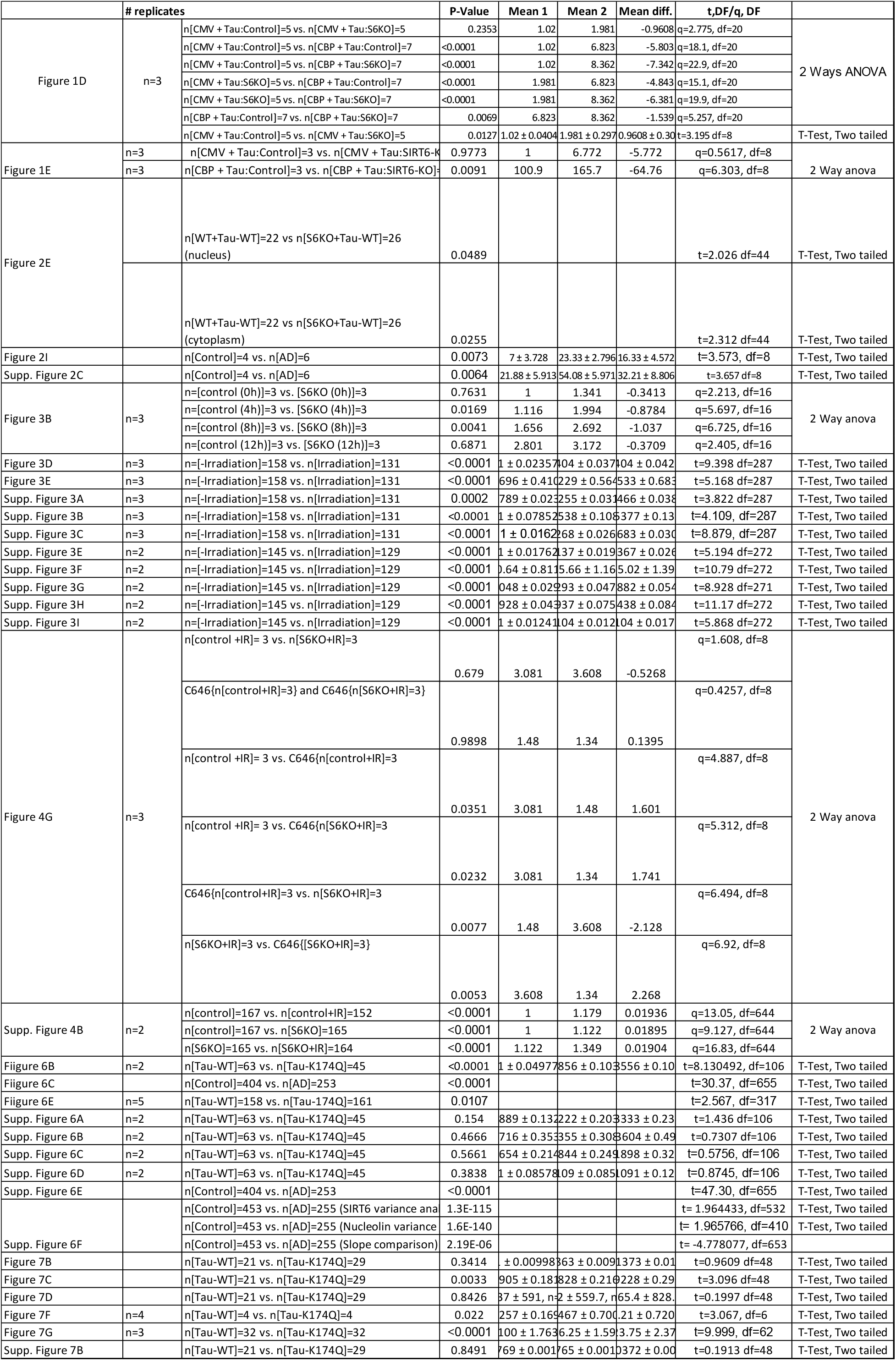

